# Lineage-specific, fast-evolving GATA-like gene regulates zygotic gene activation to promote endoderm specification and pattern formation in the Theridiidae spider

**DOI:** 10.1101/2022.06.10.495620

**Authors:** Sawa Iwasaki-Yokozawa, Ryota Nanjo, Yasuko Akiyama-Oda, Hiroki Oda

## Abstract

**Background:** The process of early development varies across the species-rich phylum Arthropoda. Owing to the limited research strategies for dissecting lineage-specific processes of development in arthropods, little is known about the variations in early arthropod development at molecular resolution. The Theridiidae spider, *Parasteatoda tepidariorum,* has its genome sequenced and could potentially to contribute to dissecting early embryonic processes.

**Results:** We present genome-wide identification of candidate genes that exhibit locally restricted expression in germ-disc forming stage embryos of *P. tepidariorum*, based on comparative transcriptomes of isolated cells from different regions of the embryo. A subsequent pilot screen by parental RNA interference identifies three genes required for body axis formation. One of them is a GATA-like gene that has been fast evolving after duplication and divergence from a canonical GATA family gene. This gene is designated *fuchi nashi* (*fuchi*) after its knockdown phenotypes, where the cell movement toward the formation of a germ disc was reversed. *fuchi* expression occurs in cells outside a forming germ disc and persists in the endoderm. Transcriptome and chromatin accessibility analyses of *fuchi* pRNAi embryos suggest that early *fuchi* activity regulates chromatin state and zygotic gene activation to promote endoderm specification and pattern formation. We also show that there are many uncharacterized genes regulated by *fuchi*.

**Conclusions:** Our genome-based research using an arthropod phylogenetically distant from *Drosophila* identifies a lineage-specific, fast-evolving gene with key developmental roles in one of the earliest, genome-wide regulatory events, and allows for molecular exploration of the developmental variations in early arthropod embryos.

## Background

The early processes of development and reproductive strategies of animals vary among species. In certain metazoan phyla, including Chordata and Arthropoda, a high degree of early developmental variations among species is observed, despite the identification of similar morphological traits and gene expression patterns during mid-embryogenesis in each phylum [1,2,3,4,5,6,7]. Mechanisms of animal evolution that account for varied developmental trajectories preceding the phylotypic period are poorly understood.

The species-rich phylum Arthropoda, comprising four major groups (Chelicerata, Myriapoda, Crustacea, and Hexapoda), conspicuously shows the diversification of early developmental processes without the disruption of stable embryonic traits, such as the elongating germ band and repetitive body units. Genetic mutation screens and a range of molecular and genetic techniques are feasible for the model organism *Drosophila melanogaster*, which has contributed to the identification of numerous essential genes and their interactions in body axis formation, germ layer formation, and segmentation. The *Drosophila* paradigm of genetic programs for early embryogenesis has allowed for candidate-gene approaches toward elucidating the evolution of developmental mechanisms in other insects or arthropod species. Such candidate-gene approaches, however, are biased with respect to the detection of key developmental genes; having relatively low consideration for lineage-specific, fast evolving, or orphan genes [8,9]. Conversely, comparative genomics in Arthropoda and other metazoan phyla has revealed lineage-specific genomic events, including gain/loss, expansion, or divergence of a gene or a gene family [10,11,12,13]. *bicoid, spätzle*, and *gurken* in *Drosophila*, which are key regulators in the organization of the major embryonic axes, are examples of lineage-specific or fast-evolving genes [14,15,16]. Considering these *Drosophila* cases and similar ones in different phyla [17,18,19], an unbiased way of identifying key developmental genes is required. Therefore, several studies that used non-*Drosophila* insect species have performed genetic mutation or gene knockdown screens by assessing larval cuticle phenotypes [20,21,22,23]. However, the identification and characterization of novel genes with key functions in specific processes of early embryogenesis are hardly feasible in any non-*Drosophila* arthropods, particularly in non-insect arthropods.

In arthropods, many model species have continued to emerge, with the genome sequenced [10,24]. There have been numerous technical merits with respect to the study of early embryogenesis in one of them, the common house spider*—Parasteatoda tepidariorum*, and distinct differences from *Drosophila* have been observed in the early embryogenesis mechanism [25]. These differences include early completion of cellularization [26,27] and involvement of Hedgehog signaling in establishing the polarity of the first embryonic axis and regulating symmetry-breaking movement of the Dpp signaling center during the orthogonalization of the first and second embryonic axes [28,29,30]. In the *P. tepidariorum* experimental system, genetic regulations during early embryonic processes can be investigated using a simple gene knockdown technique, that is, parental RNA interference (pRNAi) [29]. Research using *P. tepidariorum* has been empowered by genomic and bioinformatic resources [25,31,32,33,34,35,36], thus providing the foundation for identifying key genes involved in *P. tepidariorum* development.

The egg of *P. tepidariorum* is spherical in shape (Fig. 1). As the development starts, the egg nuclei synchronously divide, approaching the surface of the egg (stage 1). Cellular organization of the embryo is established by the time the nuclei increase to 16 [26]. The initial blastoderm comprises approximately 64 cells (stage 2; 11 h AEL), showing spherical symmetry at the morphological level. This symmetry is broken at approximately 15 h AEL by the emergence of uneven cell densities toward the formation of a germ disc within a hemisphere of the egg (stage 3) (Fig. 1; Additional file 1: Movie S1), which side is designated the embryonic side. At the pole on this side, a small number of cells is apically constricted, thus forming a blastopore [25,28]. Zygotic gene activities are required for the process of germ-disc formation [37]. Although most blastoderm cells participate in the formation of the germ disc, cells derived from around the abembryonic pole do not, and remain in the non-germ-disc region (Fig. 1; see also Additional file 1: Movie S1). The formed germ disc is sharply demarcated by the difference in cell density and comprises a single layer of more than 1,000 cuboidal epithelial cells [38]. Cells at the center of the germ disc, corresponding to the blastopore, internalize to become the central endoderm (cEND) cells and the cumulus mesenchymal (CM) cells [28,32]. The latter cell populations are clusterd and act as a dynamic source of Dpp signaling to promote the development of the dorsal side of the embryo [28,29] (Fig. 1). Cells at and near the rim of the germ disc internalize to become the peripheral endoderm (pEND) and mesoderm (pMES) cells [32]. A line of cells along the rim of the germ disc express specific genes, including a *P. tepidariorum hedgehog* homolog (*Pt-hh*), that contributes to the anterior-posterior patterning of the embryonic field [30,39,40]. The germ disc is transformed into a bilaterally symmetric germ band through orchestrated cell rearrangements, while being progressively patterned [29,38] (Fig. 1).

**Fig. 1.**
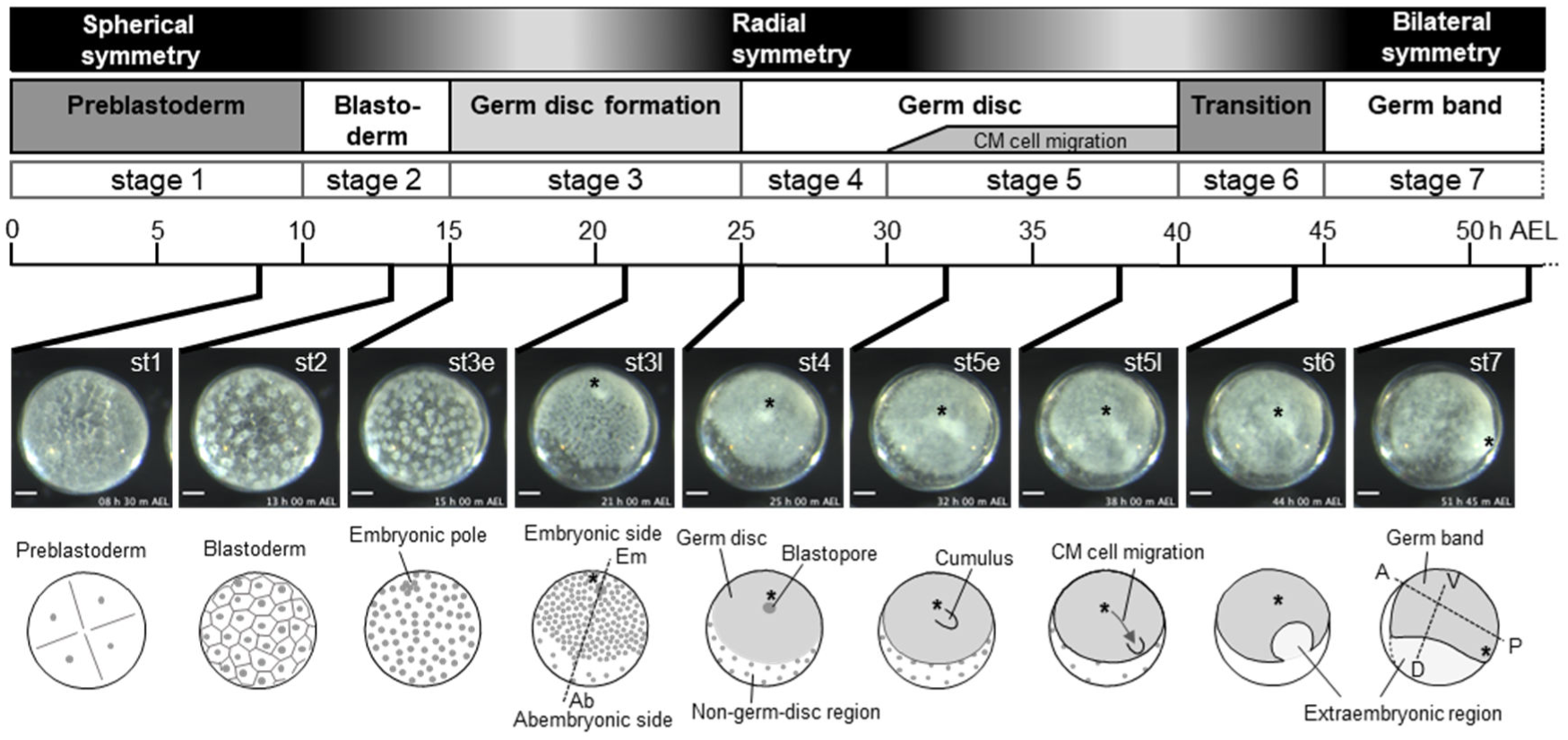
Early embryonic processes of *Parasteatoda tepidariorum*. Selected time-lapse images of a live wild-type embryo (Additional file 1: Movie S1) are shown along with their timelines (time [h] after egg laying [AEL]), stages of development, and illustrations of morphological characteristics. The morphology of the developing *P. tepidariorum* embryo undergoes symmetry transitions as shown (top). An axis of radial symmetry (embryonic-abembryonic axis) and two axes of bilateral symmetry (anterior-posterior [A-P] and dorsal-ventral [D-V] axes) are indicated by broken lines. Asterisks mark cells around the embryonic pole, cells at the closed blastopore, and cells at the posterior pole, which are in the same lineage. Scale bars,100 µm.

The formation of the sharply demarcated germ disc is an early morphogenetic event in the embryogenesis of Theridiidae and some other Araneoidea spiders [27,28,37,41,42,43,44], but it is not a general feature of spider (Araneae) embryonic development [45,46,47]. There are many other variations in early developmental processes among spider species, including the presence/absence of a visible cumulus and various modes of gastrulation [44,48]. The spider lineage provides diverse developmental trajectories prior to reaching the embryonic traits typical of arthropods.

To explore the variation in the process of early embryonic development across the phylum Arthropoda at molecular resolution, we conducted comparative transcriptomes of cells isolated from different regions of the embryo at the germ-disc forming stage in *P. tepidariorum*. Through a pRNAi-based functional screen of differentially expressed gene candidates, we identified a lineage-specific GATA-like gene, *fuchi nashi* (*fuchi*), whose knockdown hindered the completion of germ-disc formation. We obtained genome-wide datasets that represented gene expressions and chromatin accessibilities affected by *fuch*i knockdown in early embryos. Our findings showed that *fuchi* regulates zygotic gene activation to promote endoderm specification and pattern formation in the early stage of the Theridiidae spider embryo. This study provides new molecular clues for exploring the mechanisms for diversifying early developmental trajectories in the phylum Arthropoda.

## Results

### Comparative transcriptome analyses of cells isolated from different regions of early *P. tepidariorum* embryos

To search for genes with locally-restricted expression in *P. tepidariorum* embryos at stage 3, we performed RNA sequencing (RNA-seq) of small cell populations (approximately 10-30 cells) isolated from central (c), intermediate (i), and peripheral (p) regions of the nascent germ disc using glass capillary needles (Fig. 2A, B). RNA-seq reads were mapped to the *P. tepidariorum* reference genome Ptep1.0 (GCA_000365465.1), and then the reads were counted against the AUGUSTUS gene models (aug3.1) [31]. The read counts were compared using edgeR [49] by setting the following three combinations: comparison I, c cells versus i/p cells; comparison II, p cells versus c/i cells; and comparison III, i cells versus c/p cells (Fig. 2C-E). Based on these comparisons, we genome-widely identified candidates of differentially expressed genes (DEGs) (Additional file 2: Tables S1-S3). Using the values of false discovery rate (FDR) and fold change (FC), the candidate DEGs were prioritized (lower FDR with log_2_FC < -10 [for comparisons I and II] or log_2_FC > 10 [for comparison III]), selecting the top 10 genes from comparison I as Group C (Fig. 2C), the top 5 genes from comparison II as Group P (Fig. 2D), and the top 5 genes from comparison III as Group CP (Fig. 2E). The top-ranked *g4238* in Group C was listed in Group CP, despite its much stronger expression in c cells than in other cells. To avoid redundancy, *g4238* was removed from Group CP. Therefore, a total of 19 genes were systematically selected as high-priority candidate genes that might exhibit locally-restricted expression in the stage-3 embryo.

**Fig. 2.**
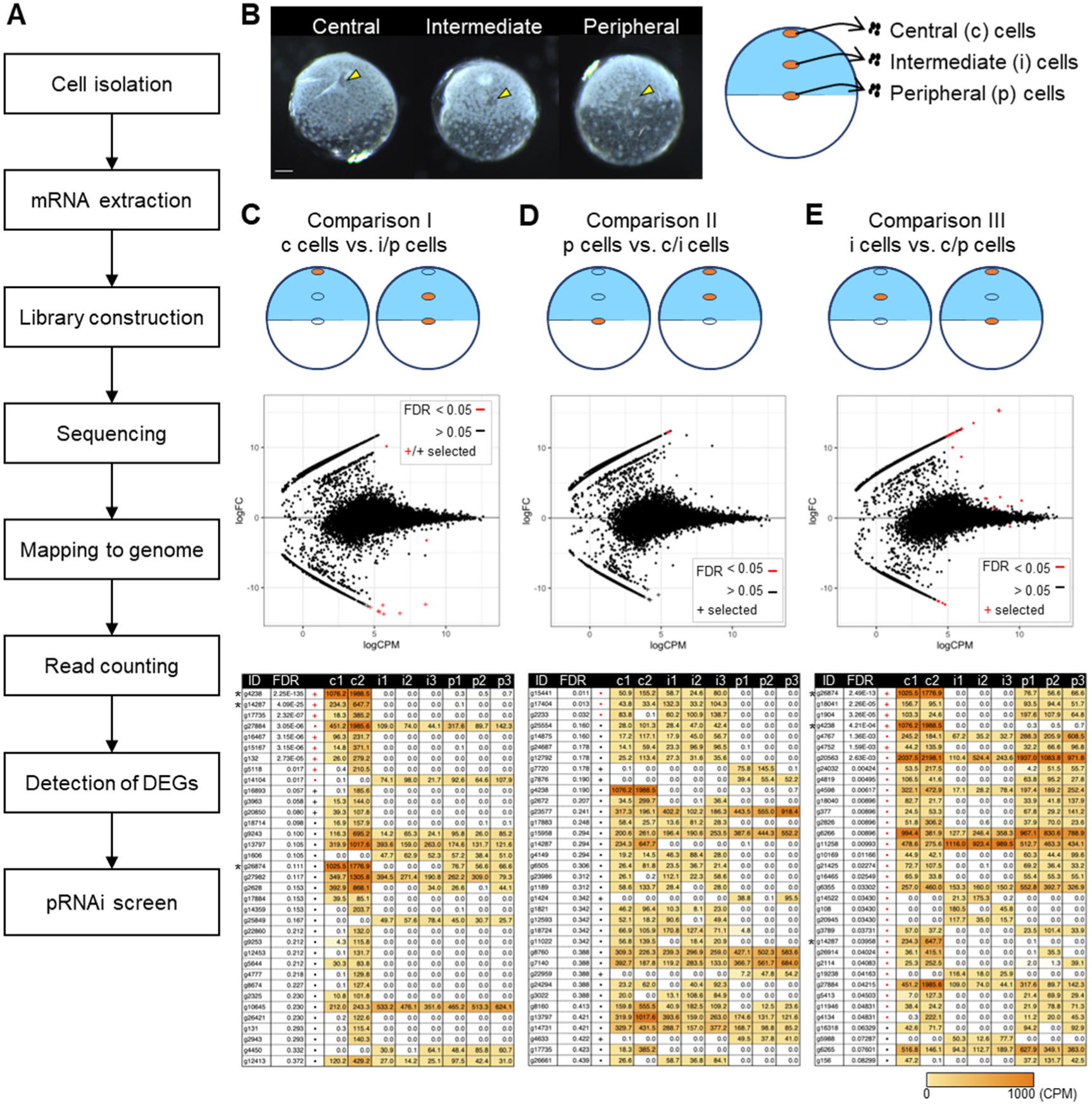
Genome-wide identification of candidate genes with locally restricted expression in stage-3 embryo. **(A)** Outline of the experimental procedure. **(B)** Images showing live stage-3 embryos following isolation of cells from 3 different regions (yellow arrowheads; central [c], intermediate [i], and peripheral [p]) of the embryo, along with the schematic showing the isolated cells used for RNA-seq. Scale bar, 100 µm. **(C–E)** Three types of comparison using RNA-seq datasets from the c, i, and p cells (comparison I: c cells versus i/p cells [C]; comparison II: p cells versus c/i cells [D]; comparison III: i cells versus c/p cells [E]) detected candidates of differentially expressed genes (DEGs). Top: schematic showing the grouping of the datasets for comparison. Middle: MA-plot of log_2_ fold-change (FC) versus log_2_ average expression level (CPM; count per million) from 10,862 genes. Bottom: lists of DEG candidates with normalized expression levels (CPM) from biological replicates of the three sample types (c1, c2, i1, i2, i3, p1, p2, and p3), which are sorted by FDR values. Genes with FDR < 0.05 are highlighted in red, and candidates of DEGs that were selected for a pilot pRNAi screen are indicated by plus signs in the MA-plots and tables. Each table displays the top 35 genes; the full lists are presented in Additional file 2: Tables S1–S3. Note that genes marked by asterisks appear in multiple tables.

To validate our gene-selection strategy, we examined the expression patterns of the 19 selected genes in stage-3 embryos using chromogenic whole-mount in situ hybridization (WISH) (Additional file 3: Fig. S1). Three of the ten Group C genes (*g4238*, *g14287*, and *g16467*) showed specific expression at the embryonic pole. Furthermore, three of the other Group C genes (*g15167*, *g5118*, *g20850*) and one of the four Group CP genes (*g26874*) showed specific expression at the embryonic pole and in a broad area on the abembryonic side. No other Group CP and Group P genes, however, showed specific detectable signals.

Additionally, similar comparative transcriptome analyses of isolated cells were applied to stage-4 and early stage-5 embryos. The resulting DEG lists (Additional file 2: Tables S4, S5) included some of the DEGs identified by the analyses of the stage-3 samples (e.g., *g4238*, and *g132*), as well as genes that had been known to show region-specific expression at the corresponding and/or later stages (eg., *Pt-lab1* [*g7954*], *Pt-BarH1* [*g8250*], *Pt-prd2* [*g18397*], and *Pt-hh* [*g4322*]).

These data suggest the effectiveness of our gene-selection strategy based on comparative transcriptome analyses of isolated cells from early *P. tepidariorum* embryos.

### Pilot functional screen identifies three genes required for germ-disc formation and/or axis formation

To identify genes with key functions in the early embryonic development of *P. tepidariorum*, we screened the 19 candidate DEGs (from the stage-3 samples) using parental RNAi (pRNAi). Double-stranded RNA (dsRNA) synthesized for each gene, as well as that for a non-spider control gene *green fluorescent protein* (*gfp*), was injected into at least two adult females. Aliquots of eggs deposited by the females were placed in oil to monitor the developmental process from early stages under the stereomicroscope. Through this screening, we identified three genes whose knockdown resulted in uncommon defects associated with germ disc formation (*g26874* and *g7720*) and/or cumulus movement (*g26874*, *g7720*, and *g4238*) (Additional file 4: Table S6)[50]. To verify the RNAi specificity, we injected two or three dsRNAs prepared from non-overlapping regions of each positive gene and confirmed that the same phenotypes were produced depending on the genes (Fig. 3A; Additional file 3: Fig. S2; Additional file 5: Movie S2)[50]. In addition, we confirmed through WISH that the transcript levels of the target genes were reduced in the corresponding pRNAi embryos (Fig. 3B).

**Fig. 3.**
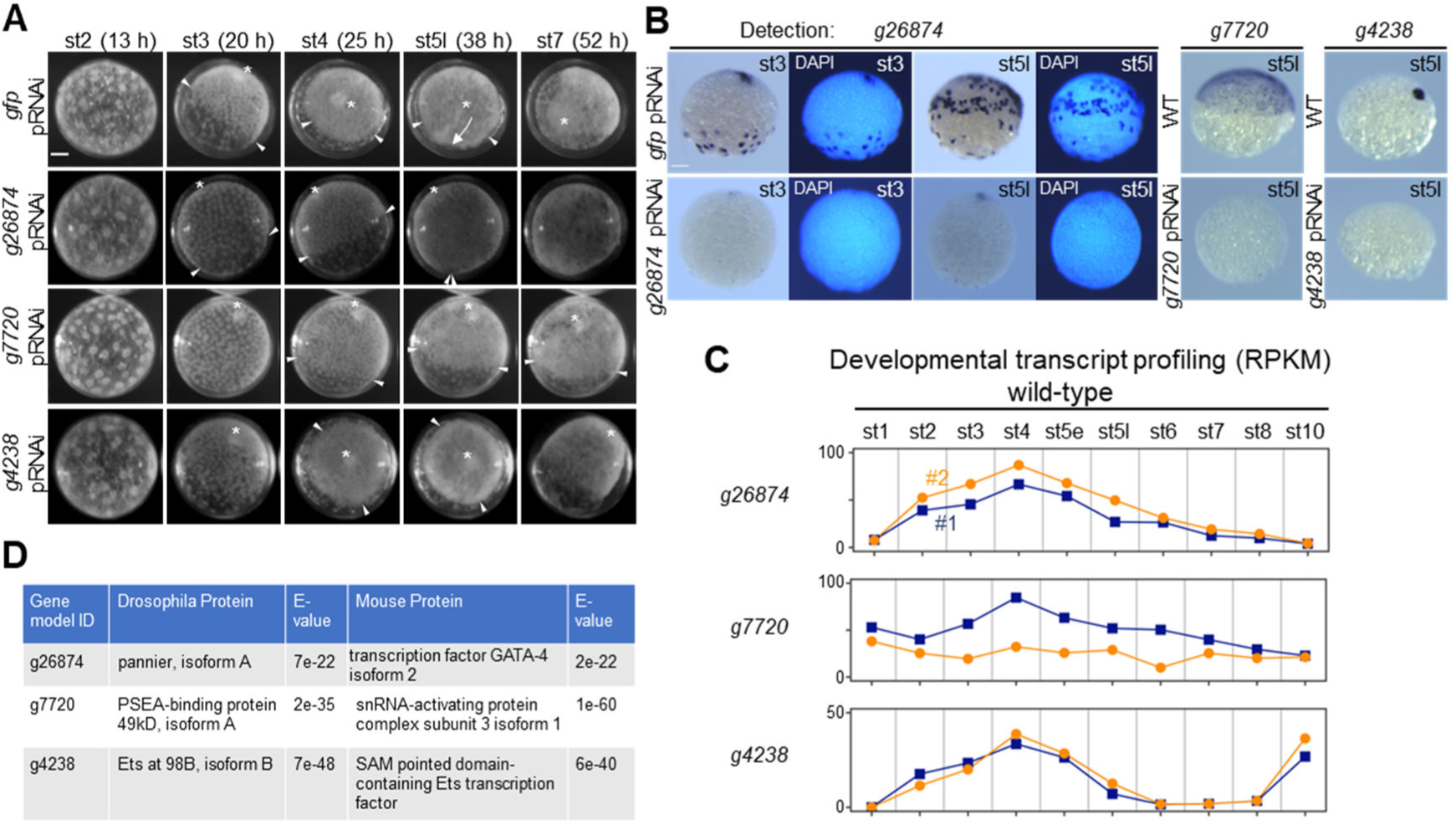
Identification of three genes in a pilot pRNAi screen of the DEG candidates. **(A)** Images showing the development of live *gfp* pRNAi, *g26874* pRNAi, *g7720* pRNAi, and *g4238* pRNAi embryos. These images are related to Movie S2 (Additional file 4). Stages of development and time (h) after egg laying (AEL) are shown. Asterisks indicate the embryonic pole, closed blastopore, and posterior pole. Arrowheads indicate the rim of the forming/formed germ disc, or corresponding sites in morphologically affected embryos. The arrow indicates CM cell migration, which did not occur in the *g26874* pRNAi, *g7720* pRNAi, and *g4238* RNAi embryos. **(B)** Effect of *g26874*, *g7720*, or *g4238* pRNAi on the expression of the corresponding target gene at stage 3 and/or late stage 5, as revealed by WISH. g*fp* pRNAi or wild-type (WT) embryos were stained as controls. Counterstains with DAPI were displayed for the comparison between *g26874* and *gfp* pRNAi embryos. **(C)** Graphs showing developmental profiling of the transcript levels (RPKM; reads per kilobase of transcript, per million mapped reads) for *g26874*, *g7720*, and *g4238* in wild-type embryos, based on public datasets [36]. Two biological replicates are individually shown. **(D)** Results of blast searches against *D. melanogaster* and mouse RefSeq protein databases using predicted amino acid sequences of *g26874*, *g7720*, and *g4238* as queries. The top-hit protein from each search is shown. Scale bars, 100 µm in A and B.

Time-lapse microscopy and cell tracking revealed that *g26874* pRNAi embryos showed normal blastoderm formation (stage 2) and subsequent initiation of cell density shift toward the formation of a germ disc, in a rather normal way, but they failed to stabilize the boundary of the forming germ disc (25 h AEL), with the cell density shift being reversed (Fig. 3A; Additional file 5: Movie S2; Additional file 6: Movie S3). In all of such severely affected *g26874* pRNAi embryos, the initial cell thickening at the embryonic pole took place. In 64% of the *g26874* pRNAi embryos (*n* = 101 from 10 egg sacs), however, CM cell migration was faint or failed to occur. The defects in germ-disc formation were not recovered at later stages, showing no clear sign of cell movements toward the formation of a germ band. In the most severe cases of *g26874* pRNAi embryos, cells that had initially shifted toward forming a germ disc reversed their movement to spread across the embryo, followed by abrupt breakdown of the surface cell layer and extrusion of the yolk materials (Additional file 5: Movie S2)[50].

*g7720* pRNAi embryos showed normal blastoderm formation followed by cell-density shift (stages 2 and 3), but its development was arrested around the end of stage 3, showing a gradually degenerating germ disc (Additional file 5: Movie S2)[50]. Although cumulus-like cell thickening was formed at the embryonic pole, it persisted there, with the overall morphology failing to develop further.

*g4238* pRNAi embryos were morphologically indistinguishable from wild-type and *gfp* pRNAi (control) embryos until early stage 5. The earliest visible difference of *g4238* pRNAi embryos from normal embryos was their failure to initiate migration of the CM cells at early stage 5, followed by their disassembly and dispersal (Additional file 5: Movie S2)[50]. In many of the embryos, surface cells around the closed blastopore formed a tail-like protrusion. Despite experiencing cumulus defects, some *g4238* pRNAi embryos displayed asymmetric cell movements toward the formation of a germ band.

### Molecular characterization of the three identified genes

Developmental transcript profiling in whole wild-type embryos, based on public RNA-seq datasets [36], revealed that there is little or no maternal supply of *g26874* and *g4238* transcripts and increasing levels of the zygotic transcripts prior to peaking at stage 4 (Fig. 3C). In contrast, substantial levels of *g7720* transcript appeared to be maternally supplied, with the zygotic transcript expressed at rather uniform levels through development.

Reciprocal blastp/tblastn searches using *Drosophila* and mouse reference sequence (RefSeq) protein databases at National Center for Biotechnology Informataion (NCBI) and the *P. tepidariorum* aug3.1 transcript sequences [31] revealed that *g7720* encoded an orthologue of *Drosophila* proximal sequence element A (PSEA)-binding protein 49kD (Pbp49) and mouse small nuclear RNA-activating complex subunit 3 (Snapc3) [51], and *g4238* an orthologue of *Drosophila* Ets98B (Dets4) and mouse SAM pointed domain-containing Ets transcription factor (Fig. 3D) [52]. *g4238* was identical to *Pt-Ets4*, as reported in a previous study [53], and consistent with our observations.

In contrast to the *g4238 and g7720* cases, reciprocal blastp/tblastn searches failed to confirm the presence of *Drosophila* and mouse orthologues for *g26874*. The best-hit *Drosophila* and mouse proteins were Pannier (Pnr), a GATA transcription factor, and GATA-4, respectively (Fig. 3D), but these proteins were significantly closer to GATA family members encoded by *g8336* and several other genes in the *P. tepidariorum* genome. Based on these results, along with those obtained from comprehensive sequence analyses described below, we concluded that *g26874* encodes a novel GATA-like gene, which we named “*fuchi nashi*” (abbreviated *fuchi*) after its knockdown phenotypes. This Japanese word means rimless.

### *fuchi* is a lineage-specific, fast-evolving GATA-like gene

The family of GATA transcription factors are characterized by the possession of a specific DNA-binding domain, which comprises two GATA-type zinc finger (ZF) motifs (ZF1 and ZF2), typically CX_2_CX_17_CX_2_C, and two basic regions followed by each ZF motif (BR1 and BR2) (Fig. 4A, B)[54,55]. Fuchi had this GATA-type DNA-binding domain, although its sequence was highly divergent from those of *Drosophila* Pnr and mouse GATA-4. To comprehensively understand the diversity of GATA family proteins among metazoans and the phylogenetic origin of *fuchi*, we exhaustively collected and manually aligned amino acid sequences of GATA family proteins from *P. tepidariorum*, other spider and non-spider chelicerate species as well as from other selected arthropod and non-arthropod metazoan species using publicly available bioinformatic resources (Additional file 7: Tables S7, S8). The resulting sequence alignment revealed that the metazoan GATA family members were classified under canonical and noncanonical types. The canonical type was characterized by a BR1 of 29 amino acid residues with high sequence similarity, and the noncanonical type by a BR1 of more, or less, than 29 amino acid residues with a varying degree of sequence divergence. The canonical GATA family members included proteins encoded by five *P. tepidariorum* genes (designated Pt-GATA1 to Pt-GATA5, encoded by *g20514*, *g1834*, *g8336*, *g25261*, and *g8337*, respectively) and their counterparts from other Araneae species, as well as *Drosophila* Serpent (Srp), the only *Nematostella vectensis* GATA (Nv-GATA), two echinoderm GATAs, and mouse GATA-1 to GATA-6 (Fig. 4B-D; Additional file 7: Table S8). Phylogenetically widespread distribution of the specific sequence features suggested that the canonical GATA family members represent the ancestral state for metazoan GATA proteins.

**Fig. 4.**
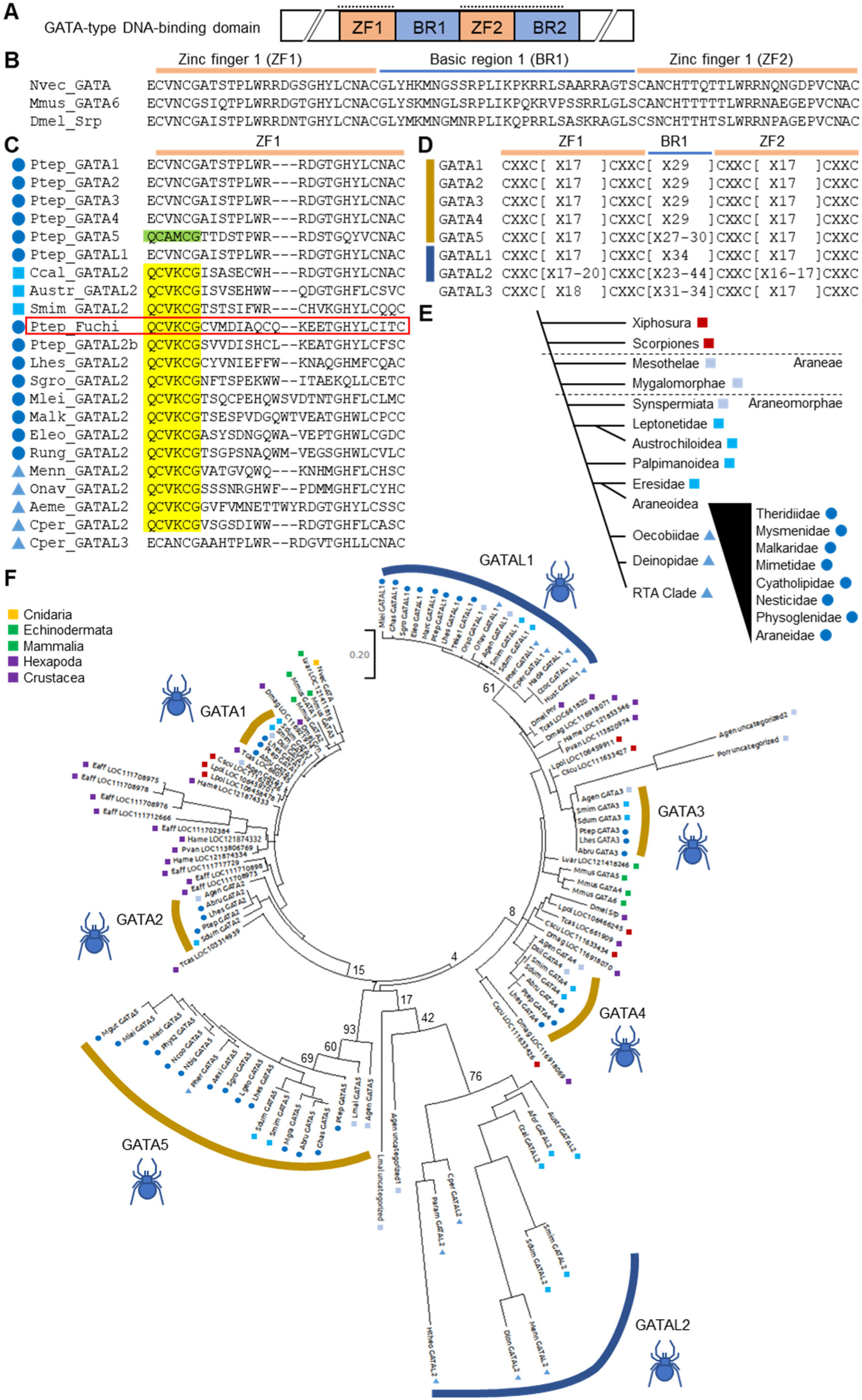
Phylogenetic characterization of Fuchi and other spider GATA family proteins. **(A)** Schematic of GATA-type DNA-binding domain. This domain comprises two zinc finger motifs (ZF1 and ZF2) and two basic regions (BR1 and BR2). **(B)** Amino acid sequence alignment of the ZF1-BR1-ZF2 region (plus one adjacent residue on the N-terminal side) of three canonical GATA family members from cnidarian, mouse and *Drosophila* (Nvec_GATA, *Nematostella vectensis* GATA; Mmus_GATA6, *Mus musculus* GATA-6; Dmel_Srp, *Drosophila melanogaster* Serpent). **(C)** Amino acid sequence alignment of the ZF1 (plus one flanking residue on the N-terminal side) of various GATA family members from *P. tepidariorum* and other spider species. There are five canonical (Ptep_GATA1 to GATA5) and three noncanonical (Ptep_GATAL1, Fuchi, GATAL2b) GATA family members in *P. tepidariorum*. The GATAL2 group, including Fuchi, is characterized by the signature sequence highlighted in yellow, and the GATA5 group by the signature sequence highlighted in green. There are two types of GATAL2 proteins; the ZF1 of one type is aligned with that of the canonical GATA without gap but the ZF1 of the other type is not. **(D)** Classification of spider GATAs based on cysteine residue spacing patterns in the ZF1-BR1-ZF2 region. GATA5s in some spider species had a BR1 of slightly varied length. **(E)** Diagram showing phylogeny of spider taxa, based on a recent study [98]. **(F)** Maximum likelihood tree of canonical and non-canonical GATA family members from Araneae, non-Araneae chelicerates, non-chelicerate arthropods, non-arthropod bilaterians, and cnidarian, using 75 amino acid sites unambiguously alignable. GATAL2 proteins with imperfectly aligned ZF1, including Fuchi, were excluded from this analysis. Bootstrap values (100 replicates) are presented at selected nodes. Color geometric codes indicate taxonomic groups of the sequence source. Details of species names and sequences are available in Tables S7 and S8 (Additional file 7).

The sequence alignment of the noncanonical GATA family members, including Fuchi, guided the identification of at least three groups specifically found within the Araneae lineage, based on gap insertion patterns, cysteine residue spacing, and sequence signatures (Fig. 4C, D; Additional file 7: Table S8). These three groups were designated as GATA-like 1 to 3 (GATAL1, GATAL2, and GATAL3). GATAL1, which included a *P. tepidariorum* protein encoded by *g26871* (Pt-GATAL1), was relatively close to the canonical GATA family members at the sequence level, but they had a BR1 of 34 amino acid residues. GATAL2, which included Fuchi and another *P. tepidariorum* protein encoded by *g26875* (designated Pt-GATAL2b), was specifically identified by a signature sequence QCV(K/R)CG at the N-terminal end of ZF1, varied cysteine residue spacing in ZF1 (CX_2_CX_17-20_CX_2_C), and a highly varied number (23-44) of amino acid residues in BR1. GATAL3, which was only found in spiders of RTA clade, was specifically identified by conserved unique sequences and cysteine residue spacing (CX_2_CX_18_CX_2_C) in ZF1. The signature sequence unique to GATAL2, QCV(K/R)CG, corresponded to ECVNCG, conserved in most canonical GATA family members (including Pt-GATA1 to 4) and GATAL1, to QC(A/V)(M/V)CG, conserved in Pt-GATA5 and its counterparts, and to ECANCG, conserved in GATAL3.

To investigate the phylogenetic relationship between canonical and noncanonical GATA family proteins, we performed maximum likelihood (ML) analysis using unambiguously aligned 75 amino acid sites from a total of 118 proteins with both ZF1 and ZF2 representing the typical cysteine residue spacing (CX_2_CX_17_CX_2_C) (Fig. 4E, F; Additional file 7: Table S8). Results showed Araneae-specific expansions of GATA protein family members, through gene duplication and divergence. ML tree topology supported the orthology of each of the GATA5, GATAL1, and GATAL2 groups. Furthermore, branch lengths indicated that the sequences of the GATAL2 proteins had been evolving much faster than those of the canonical GATA family and GATAL1 proteins (Fig 4F). Owing to their atypical cysteine residue spacing in ZF1 (CX_2_CX_18-20_CX_2_C), Fuchi and other Araneoidea GATAL2 proteins could not be included in the ML analysis (Fig. 4C; Additional file 7: Table S8). Importantly, GATAL2 proteins with the typical ZF1 (CX_2_CX_17_CX_2_C), which were used in the ML analysis, were present in a broader phylogenetic range (including Leptonetidae, Eresidae, and RTA clade) (Fig. 4C, E, F; Supplementary Table 4). This observation suggested that variously divergent sequences in ZF1 of the Araneoidea GATAL2 proteins, except the specific signature sequence, were likely due to rapid sequence evolution from the GATAL2 ancestral state.

Unlike the other two of the three genes judged as positive in the functional screen, *fuchi* was characterized as a lineage-specific gene. We therefore decided to focus our later analyses on this gene.

### *fuchi* is expressed in endoderm cells originating from both sides of the early embryo

Among the eight GATA family genes in *P. tepidariorum*, *fuchi* was the only gene which was transcribed at substantial levels during the germ-disc forming stage (stage 3) (Fig. 3C; Additional file 3: Fig. S3). We investigated the expression of *fuchi* transcript in early embryos at cellular resolution, using conventional WISH and the combination of FISH with antibody staining for β-catenin, which serves as a marker for regions of cell-cell contact (Fig. 5). Owing to technical limitations in the fixation of stage-1 embryos, the earliest stage of development examined was stage 2, the blastoderm stage (13 h AEL), when there is no morphological sign of asymmetry reflecting the future embryonic axes. At this stage, localized signals for *fuchi* expression were detected on one side of the embryo (Fig. 5A, A’, B, B’) but could not be related with the axis that would emerge a little later (15 h AEL).

**Fig. 5.**
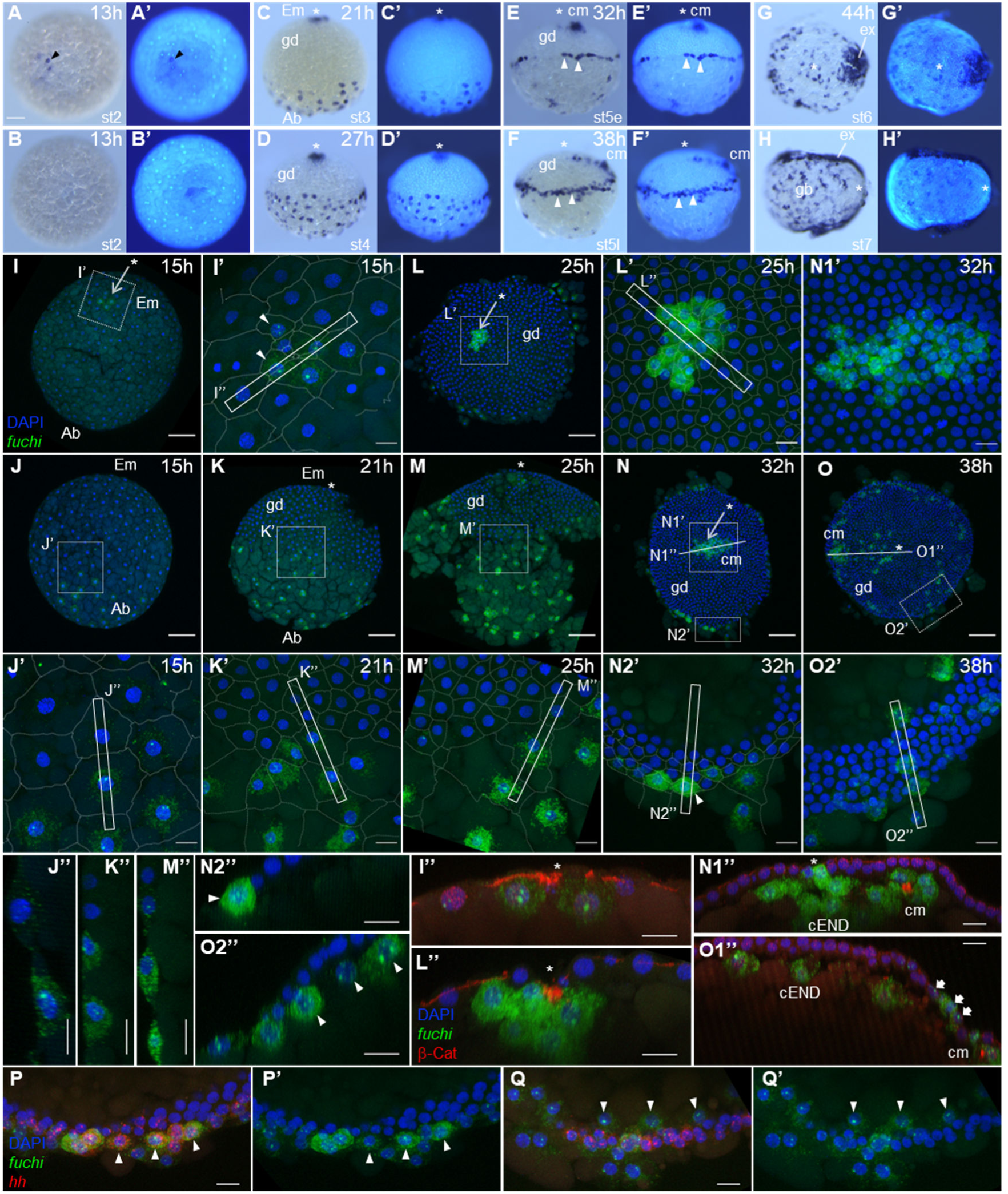
Expression of *fuchi* transcript in specific cells on the embryonic and abembryonic sides of early embryos. **(A–H)** Chromogenic WISH of wild-type embryos at indicated stages using probes for *fuchi* transcript (left), counter-stained for DNA (right). **(A, B)** Opposite sides of the same embryo. Arrowhead indicates the signal. **(C–F)** Lateral view. The forming/formed germ disc (gd) is up. Arrowheads denote cells expressing *fuchi* at or near the germ-disc rim. **(G, H)** Forming germ band (gb) with expanding extraembryonic region (ex). Asterisks mark the embryonic pole, closed blastopore, or posterior pole. **(I–O)** Reconstruction of confocal stacks showing wild-type embryos fluorescently stained for *fuchi* transcript (green) and DNA (blue) at 15 (I, J), 21 (K), 25 (L, M), 32 (N), and 38 (O) h AEL. Asterisks mark the embryonic pole or closed blastopore. I and J show the same embryo from different directions. **(I’–M’, N1’, N2’, O2’)** High magnification of the boxed areas in I–O. Cell-cell junctions visualized by staining for β-catenin are traced in I’–M’, and N2’. **(I”–M”, N2”, O2”)** Optical sections corresponding to the rectangular regions in I’–M’, N2’, and O2’. Arrowheads indicate *fuchi*-expressing cells that appear to be internalizing or internalized. **(N1”, O1”)** Optical sections corresponding to the lines in N and O. Signals for β-catenin are overlaid in red in I”, L”, N1”, and O1”. β-Catenin concentrations mark the cumulus (cm). Fat arrows indicate *fuchi* expression in the future extraembryonic region. **(P, P’, Q, Q’)** Simultaneous detection of *fuchi* (green) and *Pt-hh* (red) transcripts and DNA (blue) in cells at or near the germ-disc rim at early (P, P’) and late (Q, Q’) stage 5. Arrowheads indicate cells expressing both *fuchi* and *Pt-hh* before internalization and cells expressing only *fuchi* after internalization. Em, embryonic side; Ab, abembryonic side. Scale bars, 100 µm in A, I–O; 20 µm in other panels.

At early stage 3 (15 h AEL), the embryo begins to display morphological asymmetries, with an emerging embryonic-abembryonic (Em-Ab) axis (Fig. 1). At this stage and later, the *fuchi* transcript was detected in a small number of cells condensed at the embryonic pole (Fig. 5I, I’, I’’) and in a larger number of cells showing a progressively wider distribution on the abembryonic side of the embryo (Fig. 5J, J’, J”). Using a flat-preparation of a representative early stage 3 (15 h AEL) embryo stained for *fuchi* transcript, the number of *fuchi*-expressing cells was counted. There were 8 and 20 *fuchi*-expressing cells around the embryonic pole and on the abembryonic side, respectively, among a total of approximately 300 cells (Additional file 3: Fig. S4). Notably, signals for *fuchi* transcript in the nuclei were, in many cases, observed as paired dots (Fig. 5I’, J’), indicating that the zygotic transcription was already active at early stage 3. In the equatorial area of stage 3 embryos (15 and 21 h AEL), there was a transition from cells expressing *fuchi* transcript, on the abembryonic side, to those not expressing it, on the embryonic side (Fig. 5J, J’, J’’, K, K’, K’’). In the embryo at 21 h AEL, the *fuchi*-expressing cells had larger apical surfaces than the *fuchi*-negative cells (Fig. 5K’).

In embryos at stage 4 (25 and 27 h AEL), most *fuchi*-expressing cells on the abembryonic side had further enlarged surfaces with less tensed lines of cell-cell contact, which contrasted with a tightly packed organization of *fuchi*-negative cells constituting the nascent germ disc on the embryonic side (Fig. 5D, L, M, M’, M’’). A few *fuchi*-expressing cells, however, were identified as part of the germ disc at its border (Fig. 5M’, M”). At the blastopore, cEND cells and CM cells are internalized (Fig. 1). *fuchi* expression was observed in all these internalizing and internalized cells (Fig. 5L, L’, L’’).

In embryos at stage 5 (32 and 38 h AEL), all cells internalized through the blastopore continued to express *fuchi* (Fig. 5E, E’, F, F’, N, N1’, N1”, O, O1”). Peripheral populations of *fuchi*-expressing cells appeared to be internalized a little later than the central populations. At early stage 5 (32 h AEL) *fuchi*-expressing cells on the embryo surface were found not only in the non-germ disc region but also at the rim of the germ disc (Fig. 5N, N2’, N2’’). At late stage 5 (38 h AEL), there were *fuchi*-expressing cells that were internalized/internalizing from or through the rim of the germ disc (Fig. 5F, F’, O, O2’, O2’’). Using a DNA stain probe, SPY555-DNA, we tracked some cell nuclei in the germ-disc/non-germ-disc transition area from early stage 5 to stage 8 in a live embryo (Additional file 4: Movie S4), which directly visualized two phases of cell internalization. Phase 1 of cell internalization occurred from mid to late stage 5, where cells that were not part of the germ disc were approaching the rim of the germ disc and then internalizing to below the surface cell layer of the germ disc. These were most likely to be *fuchi*-expressing cells. Phase 2 of cell internalization occurred subsequently, where cells at the rim of the germ disc were internalizing. Double staining for *fuchi* and *Pt-hh* transcripts revealed that certain cells at and near the rim of the germ disc at early stage 5 expressed both *fuchi* and *Pt-hh* (Fig. 5P, P’). Upon internalization, however, *fuchi*-expressing cells were exclusively *Pt-hh*-negative (Fig. 5Q, Q’).

To determine whether *fuchi*-expressing cells internalized from the peripheral side of the germ disc are endoderm or mesoderm, double staining of late stage 5 and stage 6 embryos for *fuchi* and *Pt-fkh* (a marker for pMES plus pEND; [56]) or *003_J01* (a marker for pMES)[57] was performed. The use of *003_J01* enabled us to identify pMES cells prior to internalization, which was not the case with *Pt-twi*, another pMES marker [32,56,58]. Observations indicated that all internalized, *fuchi*-expressing cells were *Pt-fkh*-positive and most of them were *003_J01*-negative (Additional file 3: Fig. S5). It was also indicated that *003_J01*-positive cells appeared to be internalized later than *fuchi*-expressing cells, which presumably corresponded to Phase 2 of cell internalization. *fuchi* expression in embryos at stages 6 and 7 never showed a spatially periodic pattern along the emerging body axis (Fig. 5G, G’, H, H’), contrasting with mesoderm marker expression [58]. Taken together, these data suggested that *fuchi* is expressed in endoderm cells originating from both sides of the early embryo. In addition, *fuchi* expression occurred in association with extraembryonic differentiation (Fig. 5G, G’, H, H’). This *fuchi* expression was already initiated in the germ disc at late stage 5 (Fig. 5O, O1”).

### *fuchi* knockdown prevents the demarcation of a forming germ disc

To investigate the defects following *fuchi* knockdown, we analyzed the expression of *Pt-hh*, *Pt-fkh*, and *fuchi* transcripts in wild-type and *fuchi* pRNAi embryos at stage 4 and early stage 5 at cellular resolution. In normal germ-disc development, a line of cells expressing *Pt-hh* and *Pt-fkh* was organized to demarcate the boundary of the forming germ disc during stage 4 and early stage 5 (Fig. 6A, A’, C, C’, E, E’, G, G’). In the recognizable germ-disc/non-germ-disc transition area of *fuchi* pRNAi embryos at stage 4, the initial *Pt-hh* expression was strongly suppressed, while *Pt-fkh* expression was detectably initiated in certain cells with *fuchi*-positive nuclei and their neighboring cells (Fig. 6B, B’, D, D’). At early stage 5, faint *Pt-hh* and *Pt-fkh* expression were detected in small numbers of cells, but they did not contribute to shaping the germ disc (Fig. 6F, F’, H, H’). In *fuchi* pRNAi embryos at a later stage (late stage 5), there was no clear separation between cell populations on the embryonic and abembryonic sides, with cell internalizations being highly limited (Fig. 6I, I1’, I1”, I2’, I2”). The only cells situated below the surface layer were cells at or near the embryonic pole (Fig. 6I1’, I1”), which were associated with β-catenin concentrations marking the CM cell cluster. These observations suggested that *fuchi* knockdown prevented the demarcation of a forming germ disc and interrupted the formation of the germ layers.

**Fig. 6.**
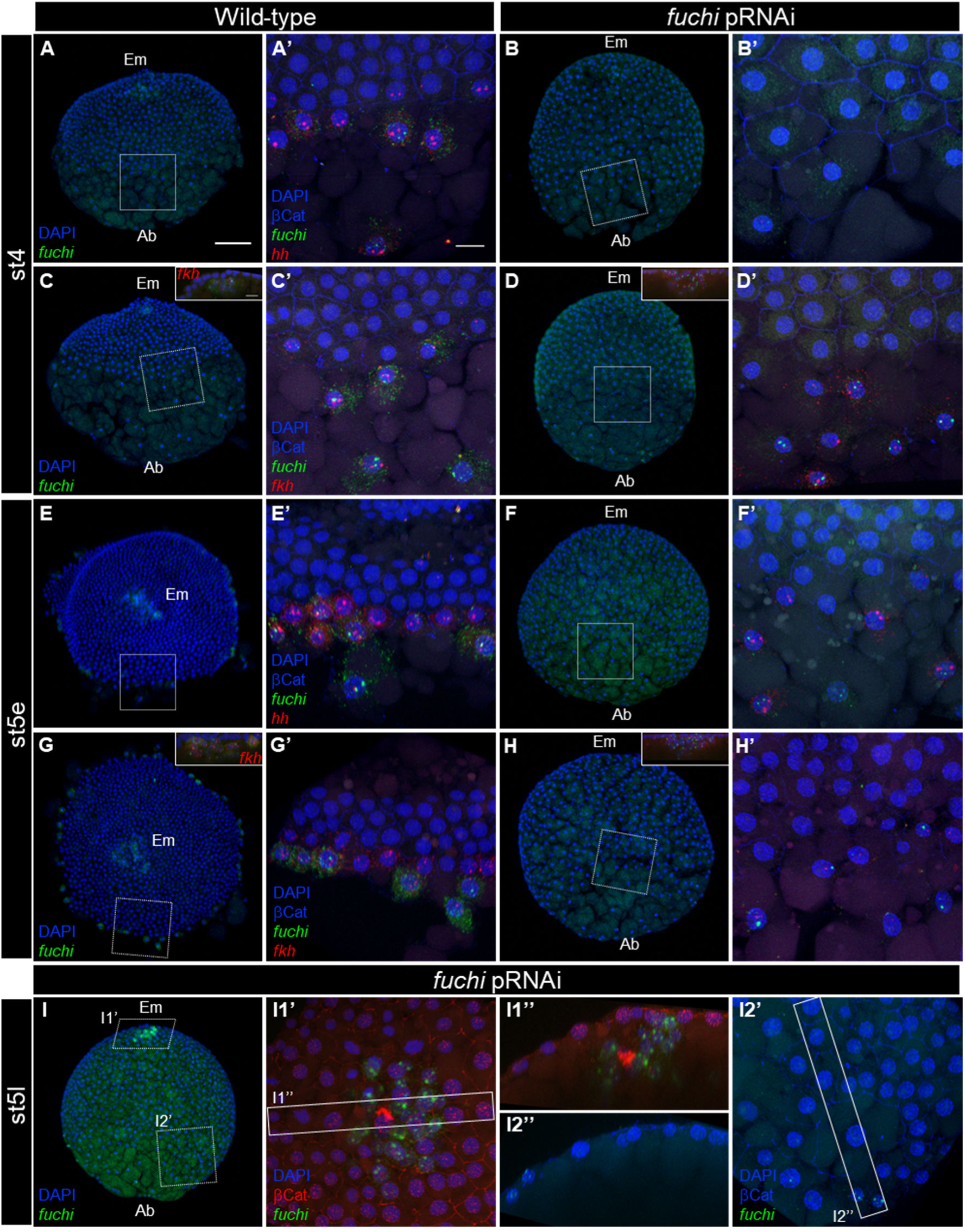
Defects of *fuchi* pRNAi embryos in demarcating a forming germ disc. **(A–H)** Reconstructions of confocal stacks showing wild-type (A, C, E, G) and *fuchi* pRNAi (B, D, F, H) embryos stained for *fuchi* (green, A–H) and *Pt-hh* (red, A, B, E, F) or *Pt-fkh* (red, C, D, G, H) transcripts, DNA (blue, A–H), and β-catenin (blue, A’–H’; not shown in A-H) at stage 4 (A–D) and early stage 5 (E–H). In A–D, F, H, and I, the embryonic side (Em) is to top, with the abembryonic (Ab) to bottom. In E and G, the embryonic side is viewed from the top. Optical sections show the presence of specific cells expressing *fuchi* at the embryonic pole (insets, C, D, G, H). The boxed areas in A-H are magnified in A’-H’, respectively. *fuchi* transcript was detectable even in cells on the abembryonic side of *fuchi* pRNAi embryos, although the signals were restricted to within the nuclei. Cells expressing *Pt-hh* and *Pt-fkh* transcripts at reduced levels were observed at regions corresponding to the the germ-disc rim and nearby abembryonic region in *fuchi* pRNAi embryos at early stage 5, but these cells did not serve as rim cells demarcating a forming germ disc. (I) Reconstructions of confocal stacks showing a *fuchi* pRNAi embryo stained for *fuchi* transcript (green, I, I1’, I2’), DNA (blue, I, I1’, I2’), and β-catenin (red, I1’; blue, I2’) at late stage 5. (I1’, I2’) High magnification of the boxed areas in I as indicated. (I1”, I2”) Cross-sections of the rectangular regions in I1’ and I2’. The cluster of CM cells were eventually internalized, but with few cells internalized in other regions. Note that the cells spread over the abembryonic side with epithelial integrity retained to a certain extent. Scale bars, 100 µm in A; 20 µm in A’, inset of C.

### *fuchi* is required for zygotic activation of endodermal and patterning genes

To explore the role of *fuchi* in regulating early *P. tepidariorum* development, we conducted comparative transcriptome analyses of *fuchi* pRNAi versus untreated embryos at stages 2, 3, and early stage 5 using RNA-seq. These analyses identified 6, 139, and 785 DEGs for stages 2, 3 and early stage 5, respectively (FDR<0.01; Fig. 7A, B; Additional file 9: Tables S9-S11). The experiments were validated by detecting lowered levels of *fuchi* transcript and unaffected levels of the α-catenin (*catA*) transcript in the *fuchi* pRNAi samples (Fig. 7C). Except for the *fuchi* gene itself, there was no overlap between the stage-2 and stage-3 DEGs but there was an overlap of 93 genes between the stage-3 and early stage-5 DEGs (Fig. 7B; Additional file 9: Tables S9-S11). According to NCBI’s gene annotation, 26 of the 93 DEGs encoded uncharacterized proteins. Transcripts for the stage-2 DEGs appeared to be maternally supplied (Additional file 3: Fig. S6), which contrasted with those for highly ranked genes from the stage-3 and early stage-5 DEGs lists (sorted by FDR value). They tended to show a sharp increase in the transcript level during stages 2 to 4, with no or little transcript supply from the mother (Fig. 7C, D). All the top 20 genes from the stage-3 DEG list were included in the stage-5 DEG list (Fig. 7D; Additional file 9: Table S10). Blastx searches against mouse and *Drosophila* RefSeq protein databases resulted in no hits for seven of the top 10 DEGs (E-value < 1e-5) (Fig. 7C). Transcripts from 8 of the top 10 DEGs (*g1125*, *g2114*, *g20850*, *g17950*, *g12453*, *g27539*, *g486*, and *g26914*) were detected in all or part of the internalized endoderm at late stage 5, and those from 3 of the 8 genes (*g17950*, *g12453*, *g20850*) were detected in cells on the abembryonic side at stage 3 (Additional file 3: Fig. S7). Notably, the *g20850* was identified as one of the Group C genes in the initial comparative transcriptome analysis of isolated cells (Fig. 2C; Additional file 3: Fig. S1).

**Fig. 7.**
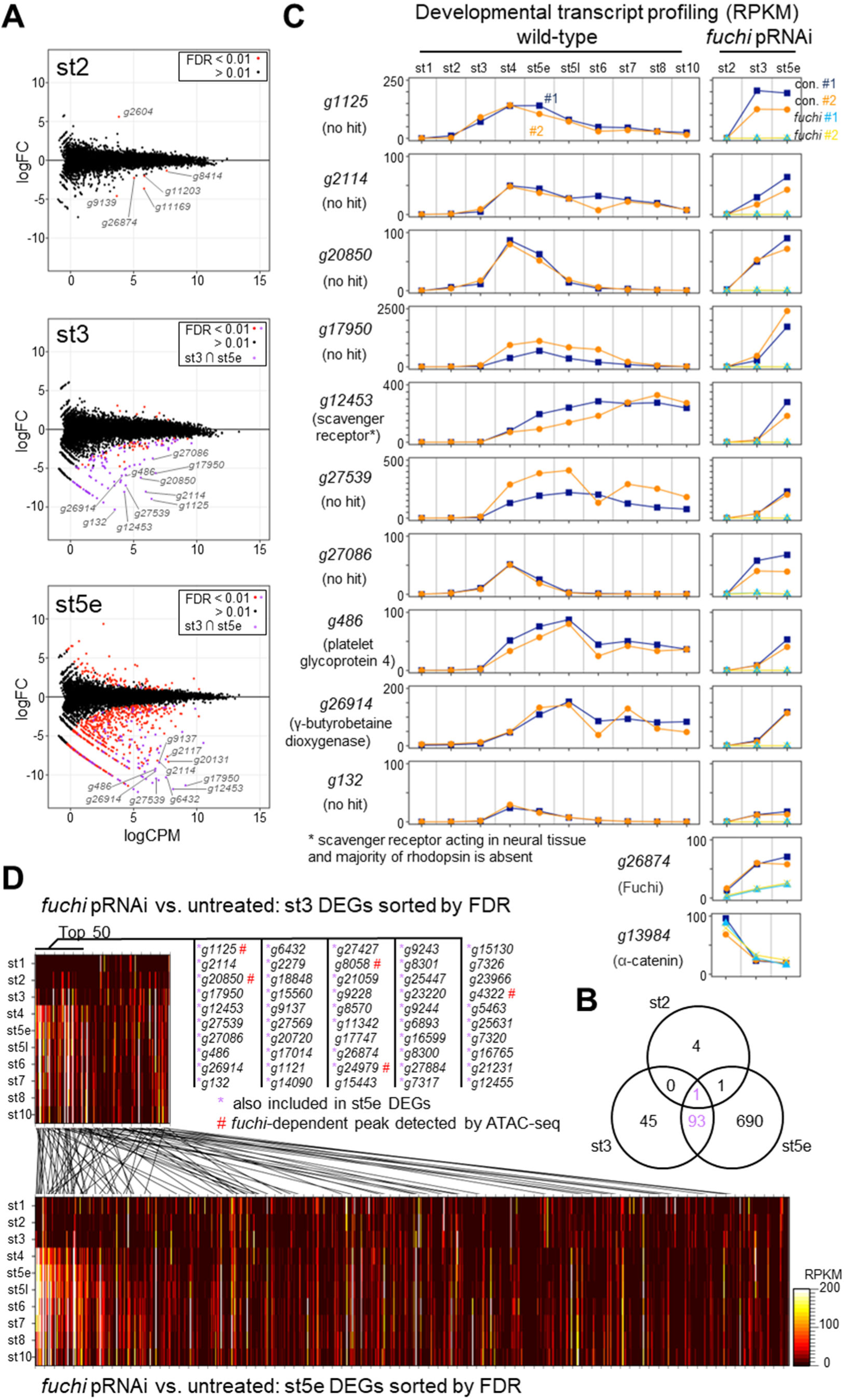
Genome-wide identification of genes whose expression levels are affected following *fuchi* knockdown. **(A)** MA-plots of log_2_ fold-change (FC) versus log_2_ average expression level (CPM) showing DEGs between *fuchi* pRNAi and control (untreated) embryos at stage 2 (st2), stage 3 (st3), and early stage 5 (st5e). Genes with FDR < 0.01 are highlighted in red or purple, with the purple genes included in both stage-3 and early stage-5 DEG lists. **(B)** Venn diagram showing the numbers of the DEGs from the three sample types and their overlaps. Note that *fuchi* (*g26874*) itself is included in all the DEG lists. **(C)** Graphs showing developmental profiling of the transcript levels for the top 10 DEGs from the stage-3 samples in wild-type embryos and the effects of *fuchi* pRNAi on the transcript levels. Top hit proteins that resulted from blastx searches against mouse and *Drosophila* RefSeq protein databases are shown, although with no hits obtained with seven of the ten genes (E-value < 1e-5). Data on *fuchi* (*g26874*) and α-catenin (*g13984*) validated the experiments. **(D)** Heat maps showing developmental profiling of the transcript levels for all the DEGs from the stage-3 (upper) and early stage-5 (lower) samples in wild-type embryos (sorted by FDR values). The top 50 DEGs from stage-3 are listed, where the genes included in stage-5 DEGs are indicated by purple asterisks. Genomic regions differentially accessible between wild-type and *fuchi* pRNAi embryos at stage 3 were found to be located within or close to some of the DEGs marked by ”#”(related to Fig. 8 and Additional file 10: Table S13).

The stage-3 *fuchi* pRNAi DEG list also included several genes previously characterized as key regulators of embryonic patterning in *P. tepidariorum*, such as *Pt-hh* (*g4322*)[30], *Pt-sog* (*g23966*)[29], and *Pt-Delta* (*g25248*)[32] (Additional file 3: Fig. S8; Additional file 9: Table S10). Despite the effects of *fuchi* knockdown on the expression of these patterning genes, certain patterning events involving *Pt-msx1* and *Pt-Delta* expression were initiated around the blastopore in *fuchi* pRNAi embryos (Additional file 3: Fig. S8). *fuchi* knockdown led to a lack of circular patterns of *Pt-hh* and *Pt-otd* expression associated with anterior patterning, but its effects on the *Pt-hh* and *Pt-otd* expression levels in whole embryos at early stage 5 were limited.

Taken together, these results suggested that *fuchi* is required for the zygotic activation of a set of endodermal genes and of some patterning genes at stage 3 or shortly before. In addition, *Pt-GATA4* (*g25261*), the likely counterpart of *Drosophila srp* (Feitosa et al., 2017), and *Pt-GATA5* (*g8337*) were included in the stage-5 *fuchi* pRNAi DEG list but not in the stage-3 one (Additional file 9: Tables S9, S10).

### *fuchi* is involved in regulating chromatin accessibility in specific genomic regions in early embryos

Certain GATA family members act as pioneer transcription factors that can influence chromatin structure [59]. Therefore, we considered the possibility that *fuchi* controls the transcription of downstream genes through chromatin structure regulation. To investigate this possibility, we conducted assay for transposase-accessible chromatin using sequencing (ATAC-seq) of wild-type (untreated), *fuchi* pRNAi, and *Pt-hh* pRNAi embryos at stage 3 (18h AEL), obtaining genome-wide datasets on chromatin accessibility. Read counts in ATAC-seq peaks were subjected to comparative analyses, and 317 genomic regions that were differentially accessible between the *fuchi* pRNAi and untreated samples but only 3 between the *Pt-hh* pRNAi and untreated samples were identified (Fig. 8A; Additional file 10; Tables S12, S13; Additional file 11; Tables S14, S15; FDR<0.05). This indicated that the *Pt-hh* pRNAi samples served as a negative control for assessing the impact of *fuchi* knockdown. The two comparisons revealed only one shared genomic region (Fig. 8A; Additional file 10; Table S13; Additional file 11; Table S15), which potentially reflected a nonspecific effect stemming from pRNAi treatment. Therefore, 316 genomic regions where chromatin accessibility was significantly affected following *fuchi* knockdown in stage-3 embryos were identified (Fig. 8B).

**Fig. 8.**
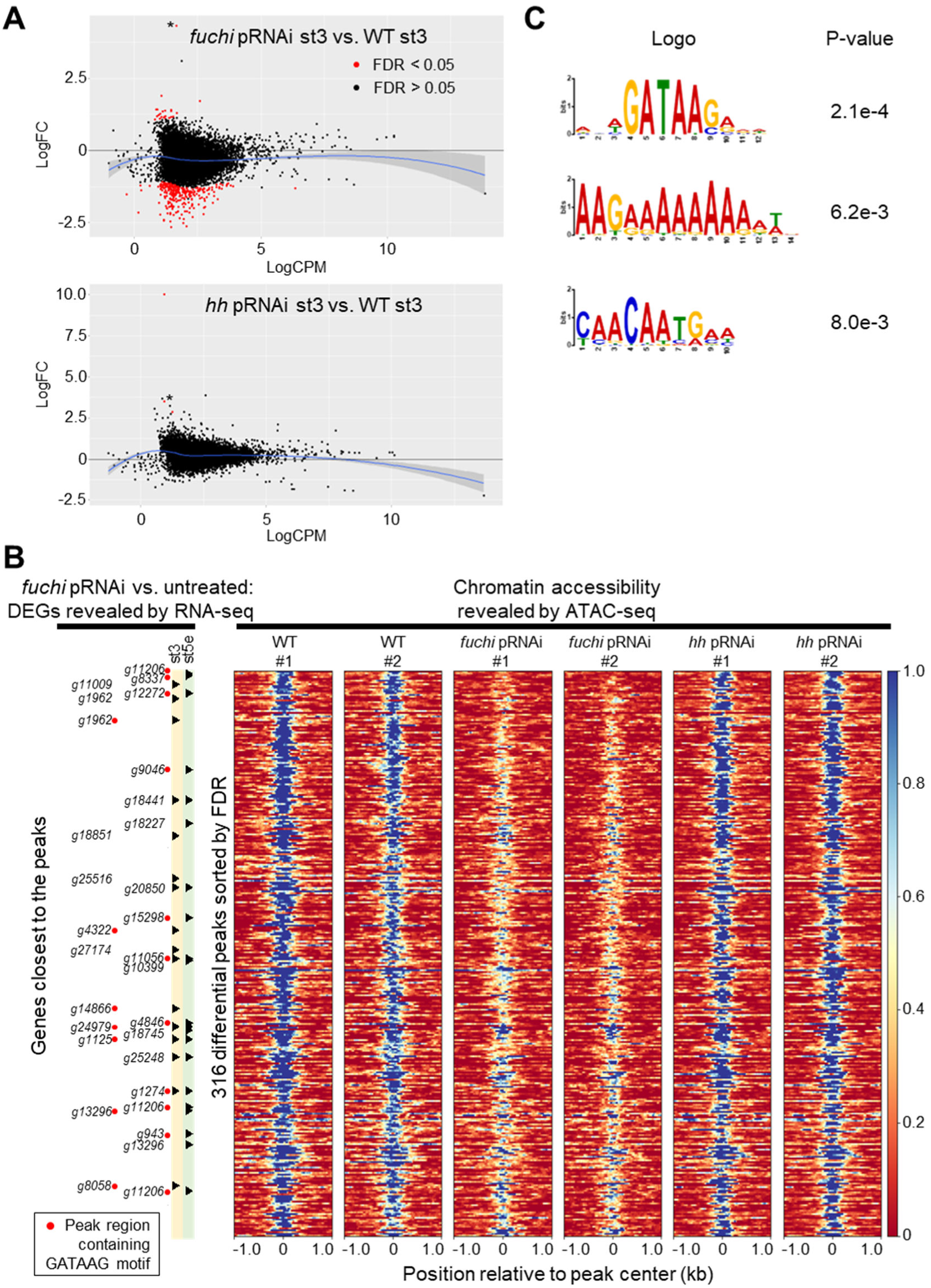
Effect of *fuchi* knockdown on chromatin accessibility in specific genomic regions in stage-3 embryos. **(A)** Distributions of differentially accessible genomic regions identified by comparisons between ATAC-seq datasets from *fuchi* (upper) or *Pt-hh* (lower) pRNAi and wild-type embryos. MA-plots represent log_2_ fold-change (FC) versus log_2_ average ATAC signal abundance (CPM). Two biological replicates were obtained from each sample type and analyzed. Significant differentially accessible regions (FDR < 0.05) are highlighted in red. Blue lines are loess fits to each distribution with 95% confidence intervals shaded in gray. Asterisks indicate positive peaks at the same genomic region. Detection of this region might not be specific to *fuchi* activity. **(B)** Heat maps showing chromatin accessibilities in the 316 genomic regions identified as *fuchi*-dependent (sorted by FDR values). Data from the biological replicates are individually shown. Some of the identified regions are located close to or within the loci of DEGs identified by the RNA-seq analyses, as shown on the left side. **(C)** Motif sequences enriched in the *fuchi-*dependent peaks, as revealed by a motif discovery tool STREME [97]. Peak regions containing detected motif sequences, RNWGATAAGAVW, are indicated in B (red dots).

Most of these regions (276/316, 93%) showed suppressed accessibility upon *fuchi* knockdown (Fig. 8A, B), and 30 of them were found to be located closer to (or within) genes included in the *fuchi* pRNAi DEG lists (for stage-3 and/or early stage-5) than to any other annotated genes (Fig. 8B; Additional file 10: Table S13). Moreover, motif discovery search revealed that motif sequences RNWGATAAGAVW, similar to typical GATA-binding motif sequences, were significantly enriched among the sequences of the *fuchi*-dependent chromatin accessibility peaks (Fig. 8C). *fuchi*-dependent chromatin accessibility peaks containing these motif sequences were actually present within or close to some of the genes identified by the comparative transcriptome analyses of *fuchi* pRNAi versus untreated embryos (Fig. 8B).

Notable examples of such genes were the top-ranked gene from the stage-3 *fuchi* pRNAi DEG list (*g1125*), the key patterning gene *Pt-hh* (*g4322*), and the canonical GATA family member *Pt-GATA5* (*g8337*) (Fig. 9A, B, C). *g1125* was predicted to encode an uncharacterized product containing two potential transmembrane domains (XP_015915605.1; Fig. 9D). Genes encoding homologous proteins to this product were detected in the genomes of at least three other Araneoidea species, *Araneus ventricosus* (GBO06802.1), *Argiope bruennichi* (KAF8764632.1), and *Oedothorax gibbosus* (KAG8179838.1)(Fig. 9D), but any non-Araneoidea protein sequences that showed similarity to these products (E-value < 1) were not found in the NCBI non-redundant protein sequences collection (March 7, 2022). *Pt-GATA5* transcript, like *fuchi*, was observed in cEND, CM, and pEND cells at stage 5 and widely distributed endoderm cells at stage 7 (Fig. 9E, F). Altogether, these results suggested that *fuchi* is required for establishing “open” states of chromatin structure in specific genomic regions, some of which are associated with endodermal and patterning genes expressed in the early stages.

**Fig. 9.**
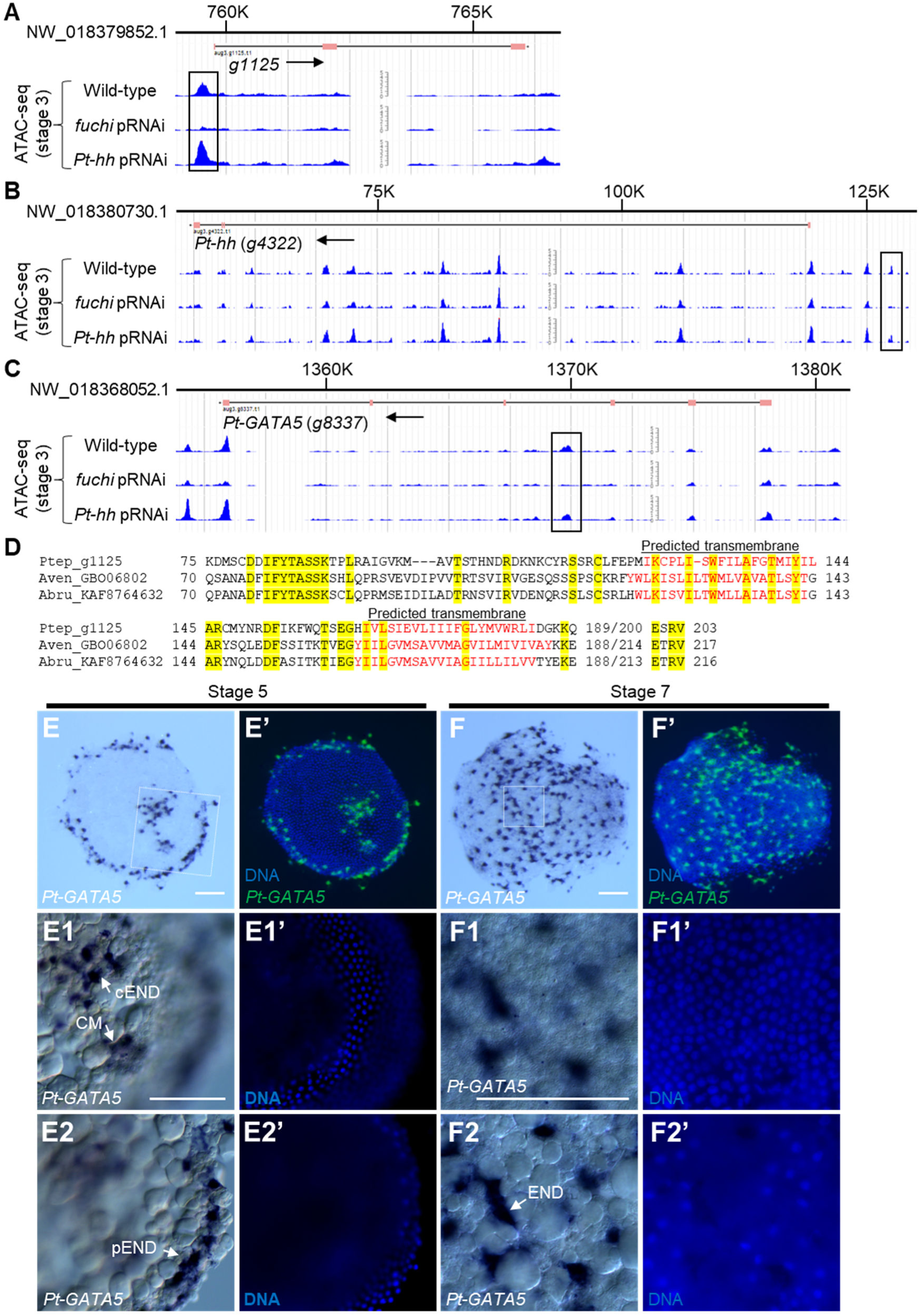
Three notable examples of genes revealed by combined analyses of the RNA-seq and ATAC-seq datasets. **(A–C)** Normalized ATAC-seq signal of stage-3 wild-type, *fuchi* pRNAi, and *Pt-hh* pRNAi embryo at the *g1125* (A), *Pt-hh* (*g4322*) (B), and *Pt-GATA5* (*g8337*) (C) loci. Line with ticks at the top represents the scaffold sequence, below which annotated exons (pink boxes) and introns (black lines) are shown. *fuchi*-dependent peak regions are indicated by open rectangles. Arrows indicate the orientation of the gene. **(D)** Amino acid sequence alignment of the *g1125* protein and its homologs detected in the genomes of two other spider species *Araneus ventricosus* (GBO06802.1) and *Argiope bruennichi* (KAF8764632.1). Residues identical across all proteins are presented with a yellow background. Residues shown in red indicate regions predicted as transmembrane domains by InterProScan version 5.53-87.0 [99]. **(E, F)** Detection of *Pt-GATA5* transcript and DNA (E’, F’) in stage-5 (E) and stage-7 (F) embryos using chromogenic WISH. **(E1, E2, F1, F2)** High magnification of the boxed areas in E and F, along with counter staining for DNA (E1’, E2’, F1’, F2’). The focal plane is at two different depths; the plane shown in E2 and F2 is deeper than that in E1 and F1, respectively. Signals are observed in cEND, CM, and pEND cells, which are located below the germ-disc epithelial layer, and in evenly distributed endoderm (END) cells, which are located below the germ-band epithelial layer. Scale bars, 100 µm.

### Detection of consistent endodermal gene expression based on single-nucleus and single-cell RNA-seq datasets

In a recent separate study, we performed single-nucleus and single-cell RNA-seq analysis of late stage 5 embryos in *P. tepidariorum* [60]. In this study, clearly isolated endodermal cell clusters, which were marked by a known early endoderm marker 012_A08 [56], were identified (Additional file3: Fig. S9). Strong *fuchi* expression was detected in these clusters, with weaker levels of *fuchi* expression being additionally detected in mesodermal (marked by *Pt-twi*)[58] and dorsal germ-disc (marked by *Pt-GATA1*)[60] cell populations. At the same time, the endoderm-specific expression of *Pt-GATA4* and *Pt-GATA5* was confirmed. In contrast to *fuchi*, and the two canonical GATA genes, *Pt-hh* expression was missing or largely reduced in the endodermal cell populations at late stage 5. These data were consistent with the staining data shown in Fig. 5 and Fig. S5 (Additional file 3). The single-nucleus and single-cell RNA-seq datasets further confirmed that most of the high-ranking genes from stage-3 *fuchi* pRNAi DEG list showed endoderm-specific expression at late stage 5 (Additional file 3: Fig. S10).

## Discussion

We conducted genome-based research using the spider *P. tepidariorum* to explore the variations in the mechanisms of early embryonic development in the phylum Arthropoda. A key initial attempt in this study was to analyze transcriptomes of cells isolated from different regions of the early spider embryo. This allowed for the genome-wide identification of candidate genes whose expression might be locally restricted along the emerging first embryonic axis, without relying on knowledge from other organisms. Although similar approaches to identifying localized transcripts in early animal embryos have been reported in non-arthropod animals [17,61,62], early arthropod embryos are rarely studied in such a way.

A benefit of using *P. tepidariorum* is the applicability of a simple gene-knockdown technique, that is, pRNAi. This allowed us to carry out a functional screen of some of the listed genes. Gene-specific embryonic phenotypes were obtained using 3 of the 19 genes tested (15%). Compared with similar screening efforts we previously made following sequence-based gene selection, this small proportion is not surprising [30]. Although similar RNAi-based gene knockdown is available in certain arthropods, such as *Tribolium castaneum*, it is important to note that the *P. tepidariorum* system allows for the easy monitoring of early embryonic developmental processes, even following pRNAi treatment. Time-lapse recording of the phenotypes from early stages gave sharable, rich, objective information on gene functions and the timing of their actions. Early defects often lead to reduced characteristic morphologies at later stages. Exclusion of common spontaneous defects is required for effective functional screening. Direct detection of the earliest developmental defects following knockdown treatments in the initial screening could simplify the structuring of our interests. Despite the small scale of the functional screen we conducted, incomparable with the large-scale, unbiased RNAi screen in *Tribolium* [23], our results may encourage additional gene-function screens in *P. tepidariorum*.

Unexpectedly, our pRNAi screen found that the cell movement toward the formation of a germ disc was reversed following the knockdown of a GATA-like transcription factor gene, which we named *fuchi nashi* (*fuchi*). This phenotype suggests the presence of a shape-stabilizing phase in the germ-disc development, which likely corresponds to stage 4, during which a line of cells along the border is organized to gain specific gene expressions. *fuchi* expression, however, is initiated much earlier than this stage, in cells outside a forming germ disc, and later continues in all internalizing and internalized endoderm cells. Our comparative transcriptome and chromatin accessibility analyses of *fuchi* pRNAi versus untreated embryos suggested that early *fuchi* activities have broad impacts on zygotic gene activation around stage 3. Lack of the *fuchi* activities appeared to hinder the specification of endoderm fates. The abnormal reversed cell movements in *fuchi* pRNAi embryos could be, at least in part, explained if we assume that the different cell types being specified exhibit different mechanical properties. The earliest morphological defects of *fuchi* pRNAi embryos possibly result from a failure to establish the differences in adhesion tension between cell populations.

We showed that early *fuchi* activities influence the expression of some of the key signaling genes involved in embryonic development and pattern formation. These include *Pt-hh*, *Pt-Delta*, and *Pt-sog*. *Pt-hh* and *Pt-Delta*, down-regulated in response to *fuchi* knockdown at stage 3, are known to show expression patterns similar to *fuchi* in early embryos (stages 3 and 4) [30,32], and *Pt-sog*, up-regulated oppositely, to be expressed in broad domains of the stage-5 germ disc [29]. Notably, early *Pt-hh* expression initiated from the abembryonic side contributes to the formation of the global polarity in the germ disc and subsequent anterior patterning. In *fuchi* pRNAi embryos, however, no patterning events that should occur from the abembryonic side of the egg were identified. These findings suggest that early *fuchi* activities specify endoderm fates and promote pattern formation in a wide area of the embryo, at the same time. Similar GATA factor-mediated regulation of *hh* expression has been documented in several mammalian and *Drosophila* developmental events [63,64,65].

The family of GATA transcription factors is known to regulate cell fate specification and cell differentiation during various developmental processes in a wide range of animals [67]. Our functional and phylogenetic characterization of *fuchi* provide the opportunity to reconsider the relationship between the evolution of GATA family members and their developmental roles. We defined “canonical” members of the GATA family, as they have perfectly alignable, close sequences to each other throughout the double zinc-finger DNA-binding domain regardless of the phylogenetic distance of the species. The canonical members include all six GATA paralogs in vertebrates, GATA-1 to GATA-6, the only *Nematostella* GATA, *Drosophila* Serpent (Srp), and five *P. tepidariorum* GATA paralogs, Pt-GATA1 to Pt-GATA5. The vertebrate GATAs are phylogenetically classified into two subgroups: GATA-1/2/3 and GATA-4/5/6, while Srp is closer to GATA-4/5/6 [55,68]. *srp* functions for endoderm specification in early *Drosophila* embryos [69], and vertebrate GATA-4/5/6 genes have developmental roles in endodermal cell lineages [70,71,72,73]. Notably, in mammals, GATA-6, in cooperation with GATA-4, is essential in the specification of the primitive and definitive endoderm, through chromatin accessibility regulation [74,75]. Based on these lines of evidence, the idea that the roles of GATA-4/5/6 genes in endoderm specification and development are conserved across bilaterians has been widely accepted. Consistent with this idea, single-cell and single-nucleus transcriptomes indicated that the likely *srp* ortholog *Pt-GATA4* (*Pt-srp*) appears to be expressed in an endoderm-specific manner at least at late stage 5 of *P. tepidariorum* development (Additional file 3: Fig. S9), although a previous report mentioned that “*Pt-srp* expression starts at stage 8 in the presumptive extraembryonic cells.” [76] Evidently, developmental transcriptomes gave a quantitative indication that there is little *Pt-GATA4* expression at stage 4 and earlier but substantial expression at late stage 5 and later (Additional file 3: Fig. S3). We suggested that instead of *Pt-GATA4*, *fuchi* and *Pt-GATA5* are key GATA family genes contributing to endoderm specification and development in the earlier stages of *P. tepidariorum* development. Although *Pt-GATA5* is a canonical member of the GATA family, it appears to have no orthologous counterpart outside the Araneae. Fuchi is not a canonical GATA factor, having a highly diverged sequence with retaining the dual zinc fingers. *fuchi* appears to act on *Pt-GATA5*, possibly through chromatin accessibility regulation. The *fuchi*-dependent ATAC-seq peak detected in the *Pt-GATA5* locus is located in an intron between exons encoding the two zinc fingers. Sequential regulation of multiple GATA factors in early endoderm specification and the subsequent differentiation process is a shared feature among distantly related bilaterians, including mouse, *C. elegans* and *Drosophila* [75,77,78]. However, the genetic components responsible for this regulation exhibit no orthologous relationships between the species, even within the Arthropoda, suggesting independent modifications of the GATA-mediated regulatory system for endoderm specification and development in the respective animal lineages.

*fuchi* and its orthologs (GATAL2) were found in sub lineages of the Araneae, including many Araneomorphae taxa but not non-Araneomorphae. These GATA-like genes, regardless of the cysteine spacing patterns in ZF1, have been rapidly evolving their sequences compared to most other GATA and GATA-like genes. Despite this divergence trend, we could define the likely orthologous GATAL2 group using the conservative signature sequence QCV(K/R)CG. Because the amino acid residues homologous to this sequence are involved in binding to specific cofactors, such as the vertebrate Friend of GATA (FOG) and *Drosophila* U-shaped (Ush), in canonical GATA factors [79,80,81], Fuchi possibly binds to a unique cofactor via this sequence. In relation with GATA cofactors, one of two *P. tepidariorum* homologs of *Drosophila* Ush (*g14866*; the other homolog is *g16893*) was included in both the gene lists obtained from the comparative RNA-seq and ATAC-seq analyses using stage-3 *fuchi* pRNAi embryos (Fig. 8; Additional file 9: Table S10; Additional file 10: Table S13). Certain GATA cofactors are known to exert an inhibitory effect to complicate the GATA factor-mediated transcriptional regulation [63,64,82]. Such an inhibitory effect of a GATA cofactor potentially accounts for the transient *Pt-hh* expression in the early abembryonic cell population. High degrees of sequence divergence in the DNA-binding domain of Fuchi and other GATAL2 proteins may be an indication of evolving interactions with cofactors and DNA-binding sites. Previous studies have suggested that N-terminal zinc fingers (ZF1) influence selectivity of DNA-binding sequences [81,83]. Alternatively, the sequence divergences might have simply resulted from reducing degrees of functional constrains. Future studies will focus on testing these possibilities.

The phylogenetic origin of the *fuchi* activity in early endoderm specification and those of the regulatory connection between *fuchi* and *Pt-GATA5*, is an intriguing issue that provides molecular clues regarding the diversification mechanism of the process of early embryonic development. The formation of a sharply demarcated germ disc is not a common feature of early spider development. In Theridiidae spider embryos, the formation of the germ disc involves an appearance of two distinct endoderm or endoderm-like cell populations derived from both polar regions of the early embryo, and the formed sharp boundary of the germ disc serves as sites of endoderm and mesoderm cell internalization, as well as sites of sending patterning signals [30,32,41,44]. These developmental strategies have potential merits of gaining parallel processes, which may allow for a more rapid development. In the embryos of many other spider species, including three distinct examples from outside the Araneoidea, *Cupiennius salei* [45], *Pholcus phalangioides* [46], and *Hasarius adansoni* [84], cell movements causing the germ layers only occur from one of the polar regions, which corresponds to the embryonic pole or the center of the germ disc in the *P. tepidariorum* embryo. The technical term “germ disc” results in potential confusions in documenting the Theridiidae embryonic development. This term is, in certain cases, used to indicate the multicell-layered region of the embryo that results from germ layer formation in spiders and other arthropods [85]. The initial germ disc in Theridiidae embryos, however, is a form of a single epithelial cell layer that precedes the formation of the germ layers [26,27,37]. *fuchi* serves as a molecular marker for the cells outside the forming germ disc. *fuchi* homologs may aid in identifying the existence of an evolutionarily conserved abembryonic cell type in other spider species. Complicatedly, the *P. tepidariorum* genome has a paralog of *fuchi* that was presumably generated by recent tandem gene duplication. No corresponding paralog was detected in other Theridiidae and non-Theridiidae spiders. The *fuchi* paralog, *Pt-GATAL2b*, shows little expression at least during the early and mid embryogenesis of *P. tepidariorum*. Studies on the expression and function of *fuchi*/*Pt-GATAL2b* and *Pt-GATA5* orthologs in the early embryos of other spider species enable us to further address the problems on the early developmental variations among the Araneae lineages.

This study has produced genome-wide datasets related to molecular and genetic regulation of early embryonic development in *P. tepidariorum*, which can be integrated with single-nucleus and single-cell RNA-seq datasets established in another study [60]. These data resources, as well as the research strategies presented, will facilitate our efforts to dissect the early developmental processes at molecular resolution and establish an independent paradigm of genetic programs for early arthropod development. Importantly, many of the genes revealed in this study encode proteins not yet characterized, thus underscoring the importance of taking genome-based approaches to studying the evolution of animal development.

## Conclusions

Using a Theridiidae spider empowered by genome sequencing, we combined comparative transcriptomes of isolated cells from different regions of early embryos and a pRNAi-based functional screen of genes. This research strategy allowed us to identify a lineage-specific, fast-evolving GATA-like gene, *fuchi*, that is essential for early embryonic development in the spider. *fuchi* is expressed in future endodermal cell poupulations in the early embryo. Further genome-wide analyses suggest that *fuchi* regulates chromatin state and zygotic gene activation to promote endoderm specification and pattern formation. The presented datasets have rich information about the molecular regulation of early embryonic development in the Theridiidae spider. The presence of many uncharacterized genes under the control of *fuchi* has been revealed. Our genome-based research using a chelicerate arthropod phylogentically distant from *Drosophila* provides the foundation for molecular and genetic exploration of the variations in early development across Arthropoda.

## Materials and Methods

### Spiders

Laboratory stocks of *P. tepidariorum* were maintained at 25 ℃ in 16 h light/8 h dark cycles. Developmental stages of the embryo were determined according to the previous descriptions [28,58]. To determine the precise time of egg laying, female behaviors were recorded every 5 min using a trail camera. The animal experimentation described was approved by the institutional committee for animal care and use (No. 2020-1) and conducted according to JT Biohistory Research Hall Regulation on Animal Experimentation.

### Cell isolation

Embryos at stage 3 were dechorionated with 50% commercial bleach for 1 min and rinsed several times with distilled water. After removing water, the embryos were placed on a glass slide with double-sticky tape and immediately covered with halocarbon oil 700 (Sigma-Aldrich). Glass capillaries (2-000-075; Drummond) were prepared in advance by pulling with a puller (PN-3; Narishige). The capillary tip was broken using forceps, just before use, and filled with lysis buffer supplied in Dynabeads mRNA DIRECT Kit (Ambion) by capillary action. Upon the manual manipulation of the capillary under the stereomicroscope, approximately 10–30 cells were sucked from a central, an intermediate, or a peripheral region of the stage-3 embryo and transferred into 0.2 ml tubes containing 10 µl of lysis buffer. The samples were stored at -80 °C until they were retrieved for RNA extraction.

### RNA-seq of isolated cells

Poly(A) mRNA was extracted using the Dynabeads mRNA DIRECT Kit (Ambion). After fragmentation of the extracted RNA for 5 min at 94 °C, first-strand cDNA synthesis and subsequent cDNA amplification by 22 cycles of polymerase chain reaction (PCR) were performed using the SMARTer stranded RNA-seq kit (Takara). The synthesized first-strand cDNA and amplified cDNA were purified using Ampure XP Beads (Beckman Coulter). Constructed libraries for sequencing were quantified using the Agilent 2100 BioAnalyzer high sensitivity DNA kit (Agilent). Two or three libraries for each region of the stage-3 embryo were sequenced in single-end runs in the antisense direction using the MiSeq reagent kit V3 (150 cycles) on the Illumina MiSeq platform. Over 10 million reads were obtained from each library. Following essentially the same procedure, RNA-seq of cells from central and peripheral regions of stage 4 and 5 embryos was performed.

### Processing of RNA-seq data

The adapter and primer sequences, as well as the first three bases derived from the SMARTer Stranded Oligo, were trimmed from the MiSeq raw reads using CLC Genomics Workbench 7.0.3 (Qiagen). Quality trimming was also performed at the following parameter settings: quality score, limit = 0.05; trim ambiguous nucleotides, maximum number of ambiguities = 2; and filter on length, discard reads below length = 30. Trimmed reads were aligned to the *Parasteatoda tepidariorum* genome (Ptep_1.0, GCA_000365465.1) using the BLAT algorithm [86] in the DNA Data Bank of Japan (DDBJ) Read Annotation Pipeline (https://p.ddbj.nig.ac.jp/pipeline/). Output alignments were filtered using the PERL script filterPSL.pl, which is accessible from the AUGUSTUS 3.0.1 scripts folder (http://github.com/Gaius-Augustus/Augustus), with the following settings: 60% coverage; 90% identity; uniqueness threshold 0.96. The filtered alignments (pslx-format) were converted to sam-format. Based on these sam-format data, the number of mapped reads was counted against the AUGUSTUS gene models (aug3.1; https://i5k.nal.usda.gov/content/data-downloads)[31]) using htseq-count v.0.6.1p1 [87] with default settings. The RNA-seq datasets, including the gene count matrix, were deposited in the NCBI Gene Expression Omnibus (GEO) database (GSE193511).

### Comparative analyses of RNA-seq gene count datasets

To assess the consistency of RNA-seq gene count datasets between biological replicates of each sample group, the coefficients of determination were calculated using the gene count matrix, and confirmed to be exceed 0.85. Out of the 27,990 annotated genes, 10,862 genes showed one or more counts per million (cpm) for at least one of the eight samples (p1-p3, i1-i3, c1, and c2). Count data matrices on these genes (10,862 genes × 8 samples) were used for comparative analyses. Differentially expressed genes (DEGs) were identified using a Bioconductor package edgeR (version 3.8.6) [49]. The following three types of comparisons were set up: comparison I, c1 and c2 vs. p1-p3 and i1-i3; comparison II, p1-p3 vs. c1, c2, and i1-i3; and comparison III, i1-i3 vs. c1, c2, and p1-p3. Two criteria were applied to determine the high-priority candidate genes: first, log_2_FC < -10 in comparisons I and II, and log_2_FC > 10 in comparison III; and second, lower FDR values.

### cDNA cloning

Full-length or partial cDNAs of newly identified genes were obtained from our collections of expressed sequence tag clones previously described [32,39], or were isolated by PCR amplification. cDNA fragments cloned in pBluescript II SK(+) (Agilent), pTriplEx2 (Takara) or pZL1 (Invitrogen) were used to synthesize dsRNA for pRNAi and/or RNA probes for in situ hybridization. The primers and cDNA clones used are summarized in Table S16 (Additional file 12).

### Parental RNAi

For dsRNA synthesis, DNA templates with the T7 promoter sequence at both ends were prepared by PCR using appropriate primers depending on the plasmid type used. dsRNAs were synthesized using a Megascript Kit (Ambion) as described previously [29]. One to two µl of the dsRNA solution at a concentration of 2.0 µg/µl was injected into the abdomen of adult females four times at 2–3 day-intervals using pulled glass capillary tubes. In the pilot pRNAi screen, dsRNA against each gene was injected into two or four females. When egg laying took place, more than 10 eggs were randomly selected from each egg sac and transferred into halocarbon oil 700, which allowed us to nondestructively monitor the embryonic development through cleared chorion.

For three genes (*g26874*, *g7720*, *g4238*) that were judged as positive in the pilot screen, 2 or 3 additional dsRNAs were prepared from non-overlapping regions of each of the cDNAs, and each dsRNA was injected into at least 2 females. Development of eggs laid by the injected females were monitored using time-lapse recording, or with occasional visual inspection, under the stereo microscope. *gfp* dsRNA was used as the control.

### Time-lapse microscopy of live embryos

Embryos dechorionated with 50 % commercial bleach were placed on a glass slide with double-sticky tape and covered with halocarbon oil 700. When the observation was started from stage 1, the dechorionation step was omitted to prevent possible effect on the development. Images were taken every 5 or 10 min using a stereo microscope (SZX12, Olympus) equipped with a color CCD camera (C7780-10, Hamamatsu Photonics) controlled by AquaCosmos software (Hamamatsu Photonics) or other stereo microscopes (M165C, Leica; or SZX7, Olympus) equipped with a color CMOS camera (WRAYCAM-G200 or -NF300, WRAYMER).

For live imaging of nuclei around the germ-disc/non-germ-disc transition area, dechorionated stage-4 embryos were placed on a glass slide with double-sticky tape, and an embryo appropriately oriented was selected and skewered with a thin glass needle to prevent the embryo from rotating during later development. A 1,000× stock solution of a cell-permeable dye SPY555-DNA (Spirochrome), which was prepared in 50 µl of dimethylsulfoxide, was microinjected into the perivitelline space of the skewered egg. The injecting volume should be low to avoid affecting the development of the embryo from being affected. Approximately 1 hour after dye injection, the embryo started to be examined under an Olympus BX50 fluorescence microscope equipped with a cooled CCD camera (CoolSNAP HQ, Roper Scientific) controlled by MetaMorph.version 6.1 (Universal Imaging). Using 10 × objective lens (UPlanAPO 10×/0.40, Olympus), bright-field and fluorescence 500 × 500 pixel images were acquired every 5 min at exposure times of 10 msec and 100 msec, respectively. To reduce the excitation light, two neutral density filters (U-ND6-2 and U-ND25-2, Olympus) were used. A single focal plane, which was manually adjusted during approximately 30 h of observation, was recorded at each time point. The observed embryo was checked for its normal development. Time-lapse images were analyzed using Imaris version 7.6.5, where cells were manually tracked, and ImageJ version 1.51d.

### Molecular phylogenetic analysis of GATA family members

To collect amino acid sequences containing GATA-type DNA binding domain from many arachnid species and other arthropod and nonarthropod metazoan representative species, we used nucleotide sequence resources presented in Table S7 (Additional file 7), which included coding sequences from sequenced genomes, transcriptome assemblies, and RNA-seq raw data, retrieved from GenBank (NCBI) or GigaScience database. RNA-seq of *Hasarius adansoni* mid-stage embryos was performed in this study, as described below. The RNA-seq data were subjected to de novo assembling using Trinity (version 2.11.0). To identify amino acid sequences coding for GATA-type DNA binding domains in the retrieved or assembled nucleotide sequences, blastp searches were performed using the Fuchi amino acid sequence as a query with a cut-off e-value of 1e-2, followed by the systematic selection of sequences that contained at least one GATA-type zinc-finger motif sequence (CX_2_CX_16-20_CX_2_C). The identified amino acid sequences were manually aligned and classified based on the detection of signature sequences, excluding GATA-like proteins that had only one GATA-type zinc finger motif. For molecular phylogenetic analysis, 75 amino acid sites from 118 sequences, which were unambiguously aligned, were used (Additional file 7: Table S8). The molecular phylogeny was inferred using the Maximum Likelihood (ML) method and JTT matrix-based model [88] on the software MEGA11 [89].

### RNA-seq of whole embryos

For RNA-seq analyses of *fuchi* pRNAi embryos, mated adult females were injected with approximately 1–2 µl of *fuchi* dsRNA solution (2 µg/µl) four times at 2- to 3-day intervals. For RNA extraction, 50–100 embryos at stage 2, 3, and early stage 5 that were derived from egg sacs produced by the injected females 2 days before (untreated) and 18–19 days after the first injection of the dsRNA were used. Two biological replicates were prepared for each sample type. The mRNA extraction and library construction for sequencing were performed as previously described [36]. The libraries were quantified using the Library quantification kit (Takara) and Thermal cycler Dice Realtime TP800 (Takara) and sequenced in single-end runs (150 cycles) in the antisense direction on the Illumina MiSeq platform. The sequence reads were processed as previously described [36], and the datasets were deposited in the GEO database (GSE193650). Estimates of DEGs between untreated and *fuchi* pRNAi embryos at each stage were performed using EdgeR (version 3.8.6).

For the *H. adansoni* RNA-seq, poly(A) mRNA was extracted using a QuickPrep Micro mRNA purification kit (GE Healthcare) from 20 sibling mid-stage embryos with extending limbs. Library construction for sequencing was performed as previously described [36]. The library was sequenced in a paired-end run (150 × 2 cycles) on the Illumina Miseq platform, and the raw reads (SRR17326784) and the *H. adansoni* transcriptome assembly (GJQJ00000000) were deposited in GenBank.

### ATAC-seq of whole embryos

ATAC-seq was performed following the procedure described by Buenrostro et al. (2015) [90] with modifications considering the procedure reported by Haines and Eisen (2018) [91]. In brief, 40 whole embryos at stage 3 (18 h AEL) were homogenized using disposable pellet pestles (12-141-368; Fisher Scientific) and 1.5 mL Eppendorf tubes (0030125150) in 50 µl lysis buffer (10mM Tris, 10mM NaCl, 3mM MgCl_2_, adjusted to pH 7.4 with HCl) containing 0.15 mM spermidine, after which 50 µl of lysis buffer and 1 µl of 10% Nonidet P-40 were added as the pestle was rinsed. After a 10 min incubation, nuclei were spun down at 800 g and resuspended in 17.5 µl water, followed by the addition of 25 µl of 2× TD buffer (Illumina) and 7.5 µl of Nextra Tn5 transposase (Illumina). Transposed DNA was purified using MinElute Reaction Cleanup Kit (Qiagen) and amplified with Illumina Nextera Transposase Adaptors index PCR primers, and the amplified DNA was purified using Ampure XP Beads (Beckman Coulter). Constructed libraries were paired-end sequenced (151 bp reads × 2) on the Illumina HiSeq X Ten system.

Comparative analyses of ATAC-seq datasets from three different sample types (wild-type, *fuchi* pRNAi, and *Pt-hh* pRNAi) were performed basically following the procedure described by Reske et al. (2020)[92] with slight modifications based on guidelines provided by Gaspar (2019)[93] and Delisle et al. (2021)[94]. Two biological replicates were obtained from each sample type. ATAC-seq raw reads were trimmed for adaptor and primer sequences and low-quality sequences using Trim Galore! with default settings. Trimmed reads were then aligned to the *P. tepidariorum* genome (Ptep_2.0) using bowtie2 v2.2.5 with default settings. Reads mapped with low quality scores (less than 30) were removed using samtools (version 1.9). PCR duplicates were removed using Picard MarkDuplicates (version 2.23.4). The alignments of filtered reads to the genome were saved as bam files. Peak calling was performed using the ATAC-seq mode of Genrich (version 6.0).

Differential accessibility analysis was performed using a Bioconductor package, *csaw* [95]. Peak regions called in both replicates of each sample type were extracted, and the peak regions extracted from the three sample types were merged. Reads were counted in peaks, with low abundance peaks (log CPM < -3) removed. Read counts were subject to normalization with the trimmed mean of *M* values (TMM) method [96]. Differential accessibility was estimated using the glmQLFit and glmQLFTest functions in EdgeR (version 3.32.0). Peaks with FDR < 0.05 were considered as significant. To discover motif sequences in the identified differential peak regions, STREME [97] was used.

Wild-type whole embryos at stages 1 (9 h AEL) and 2 (12 h AEL) were similarly analyzed by ATAC-seq to detect genomic regions differentially accessible between the different stages (stages 1 to 3), but no clear results were obtained. All the ATAC-seq datasets were deposited in the GEO database (GSE193870).

### Staining of embryos

Antisense RNA probes for in situ hybridization were prepared by *in vitro* transcription using Digoxigenin- or Dinitrophenyl-conjugated UTP (DIG, Roche 11277073910; DNP, PerkinElmer NEL555001EA) as previously described [56]. Whole-mount in situ hybridization with chromogenic substrate was performed in the previously described manner [29] with the exception of an additional tyramide-based signal amplification (TSA) process described previously [56]. In brief, after washing DIG-conjugated RNA probes, the samples were incubated with anti-DIG-POD antibody (11633716001, Roche; 1:500 dilution) and then reacted with DNP-tyramide (NEL747, PerkinElmer). In the detection step, anti-DNP-AP antibody (MB-3100, Vector; 1:50) and NBT/BCIP were used. Most samples were counterstained with 4’,6-diamidino-2-phenylindole (DAPI; Sigma-Aldrich).

Double-color FISH was performed as previously described [56]. To visualize the DIG- and DNP-conjugated probes, anti-DIG-POD (Roche 11633716001; 1:500) and anti-DNP-HRP (PerkinElmer FP1129; 1:100) were used in combination with DyLight488-tyramide and DyLight680-tyramide, respectively.

Single- and double-color FISH were combined with immunostaining for β-catenin in certain cases (Figs. 5 and 6), where Proteinase K treatment prior to hybridization was omitted. After completing the FISH procedure, the embryos were incubated with rabbit anti-β-catenin antibody (C2206, Sigma-Aldrich 1:500) and then with Cy3-conjugated anti-rabbit IgG antibody (AP187C, Chemicon; 1:200), followed by counterstaining with DAPI.

### Image acquisition and processing

Images for chromogenically stained WISH samples were obtained using a stereo microscope (SZX12, Olympus), a fluorescence unit BT-ExSMOP (Biotools), and a CCD camera (C7780-10, Hamamatsu Photonics). For confocal microscopy of fluorescently stained embryos, the whole embryos were individually mounted on glass slides with spacers of approximately 100–500 µm, which were adjusted depending on the sample using sticky tape (CT-18S, Nichiban) and silicone vacuum grease (335148, Beckman). To view different sides of a single embryo, it was rotated after being mounted on the glass slide. The embryos were examined using a Leica TCS SPE confocal system equipped with four laser sources (405, 488, 532, and 635 nm), which were used to excite DAPI, DyLight488, Cy3, and DyLight680, respectively. Acquired confocal stacks were processed and analyzed using Imaris version 7.6.5 (Bitplane) and ImageJ version 1.51d. The Imaris snapshot feature and oblique slicer tool were used to obtain images capturing the embryonic regions of interest. Linear signals of cell-cell adherens junctions visualized by β-catenin staining were traced in Adobe Photoshop CS5 ver. 12.0.4.

## Supporting information

Addfile1 Movie S1

Addfile2 Table S1-S5

Addfile3 Figure S1-S10

Addfile4 Table S6

Addfile5 Movie S2

Addfile6 Movie S3

Addfile7 Table S7-S8

Addfile8 Movie S4

Addfile9 Table S9-S11

Addfile10 Table S12-S13

Addfile11 Table S14-S15

Addfile12 Table S16

## Availability of data and materials

Raw and processed data for the comparative RNA-seq and ATAC-seq analyses are accessible from NCBI GEO (GSE193511, GSE193650, and GSE193870). The data aligned to the reference genome can be viewed on our web database (https://www.brh2.jp). The nucleotide sequences determined for *fuchi* and *Pt-GATA5* are available under the accessions LC671429 and LC671430 in the DDBJ/EMBL/GenBank International Nucleotide Sequence Databases. The sources of sequence data used in the molecular phylogenetic analysis are listed in Additional file 7: Tables S7 and S8. Supplementary movies showing the pilot pRNAi screen and the pRNAi specificity validation are available on figshare [50].

## Abbreviations

AEL: After egg laying
ATAC-seq: Assay for transposase-accessible chromatin using sequencing
BR: Basic region
CPM: Counts per million
DAPI: 4’,6-diamidino-2-phenylindole
DDBJ: DNA Data Bank of Japan
DEG: Differentially expressed gene
dsRNA: Double stranded RNA
FC: Fold change
FDR: False discovery rate
GEO: Gene Expression Omnibus
ML: Maximum likelihood
NCBI: National Center for Biotechnology Informataion
PCR: Polymerase chain reaction
pRNAi: Parental RNA interference
RNA-seq: RNA sequencing
RPKM: reads per kilobase of transcript, per million mapped reads
WISH: Whole-mount in situ hybridization
ZF: Zinc finger

## Acknowledgements

We thank Akiko Noda for technical assistance, and other members of JT Biohistory Research Hall for discussion.

## Contributions

S.I., Y.A., and H.O. designed the study. S.I. created all the RNA-seq datasets. S.I. analyzed the RNA-seq datasets with help from Y.A. S.I. performed the pRNAi screen and the gene expression and phenotype analyses. S.I. and H.O. performed the phylogenetic analysis. R.N. created the ATAC-seq datasets. R.N. analyzed the ATAC-seq datasets with help from Y.A. and H.O. S.I. wrote the first draft of the manuscript. H.O. wrote the final version of the manuscript with input from all authors.

## Ethics approval and consent to participate

Not applicable

## Consent for publication

Not applicable

## Supplementary Information

**Additional file 1: Movie S1.** Time-lapse video of sibling *Parasteatoda tepidariorum* embryos. The left embryo is the same as shown in Figure 1. Time after egg laying (AEL) is indicated. Scale bar, 100 µm.

**Additional file 2: Table S1.** List of DEG candidates identified by comparative transcriptome analysis of c versus i/p cells from stage-3 embryo (comparison I). **Table S2.** List of DEG candidates identified by comparative transcriptome analysis of p versus c/i cells from stage-3 embryo (comparison II)**. Table S3.** List of DEG candidates identified by comparative transcriptome analysis of i versus c/p cells from stage-3 embryo (comparison III)**. Table S4.** List of DEG candidates indentified by comparative transcriptome analysis of c versus p cells from stage-4 embryo**. Table S5.** List of DEG candidates identified by comparative transcriptome analysis of c versus p cells from early stage-5 embryo.

**Additional file 3: Figure S1.** Chromogenic WISH of stage-3 embryos using probes for 19 selected DEG candidates. **Figure S2.** Validation of the specificity of the RNAi effects for *g26874*, *g7720*, and *g4238*. **Figure S3.** Developmental transcript profiling of *P. tepidariorum* GATA family genes in wild-type embryos. **Figure S4.** Cell counting in a stage-3 embryo stained for *fuchi*. **Figure S5.** *fuchi* expression in pEND cells. **Figure S6.** Characterization of 5 DEG candidates from the comparative transcriptome analysis of stage-2 *fuchi* pRNAi versus untreated embryos. **Figure S7.** Characterization of the top-10 DEG candidates from the comparative transcriptome analysis of stage-3 *fuchi* pRNAi versus untreated embryos. **Figure S8.** Effects of *fuchi* RNAi on expression of selected genes. **Figure S9.** Single-nucleus transcriptome data showing endodermal and mesodermal cell populations in late stage-5 embryos. **Figure S10.** Single-nucleus and single-cell transcriptome data showing cell populations expressing the genes regulated by *fuchi*.

**Additional file 4: Table S6.** Summary of a pilot pRNAi screen of 19 selected genes.

**Additional file 5: Movie S2.** Time-lapse video showing phenotypes *gfp*, *g26874*, *g7720*, and *g4238* pRNAi embryos. Time after egg laying (AEL) and time after start of recording are indicated. Movie starts are adjusted by the timing of blastoderm formation. Some of the embryos are the same as those shown in Figure 3A. Scale bar, 100 µm.

**Additional file 6: Movie S3.** Tracking of cells in wild-type and *g26874* pRNAi embryos. In the *g26874* pRNAi embryo, cells on the abembryonic side initially shift toward forming a germ disc but, later, they return back. Time after start of recording is indicated.

**Additional file 7: Table S7.** List of bioinformatic resources used for phylogenetic characterization of Fuchi and other GATA family members in spiders. **Table S8.** Amino acid sequence alignment and classification of GATA family members from spiders and other metazoans.

**Additional file 8: Movie S4.** Time-lapse video showing cell internalizations taking place around the germ-disc rim. Nuclei labeled by SPY555-DNA are presented in white. The germ disc on the embryonic side is to the lower right, while the non-germ-disc region on the abembryonic side is to the upper left. Internalizing cells are tracked. Cells from the non-germ-disc region are moving to below the germ-disc epithelium. Later, cells at the germ-disc rim internalize. Time [d : h : m] after start of recording is indicated.

**Additional file 9: Table S9.** List of DEGs identified by comparative transcriptome analysis of *fuchi* pRNAi versus untreated embryos at stage 2. **Table S10.** List of DEGs identified by comparative transcriptome analysis of *fuchi* pRNAi versus untreated embryos at stage 3. **Table S11.** List of DEGs identified by comparative transcriptome analysis of *fuchi* pRNAi versus untreated embryos at early stage 5.

**Additional file 10: Table S12.** Data from comparative analysis of read counts in extracted ATAC-seq peaks between *fuchi* pRNAi and wild-type embryos at stage 3 using edgeR. **Table 13.** List of differential ATAC-seq peaks between *fuchi* pRNAi and wild-type embryos at stage 3 (FDR < 0.05).

**Additional file 11: Table S14.** Data from comparative analysis of read counts in extracted ATAC-seq peaks between *Pt-hh* pRNAi and wild-type embryos at stage 3 using edgeR. **Table S15.** List of differential ATAC-seq peaks between Pt-hh pRNAi and wild-type embryos at stage 3 (FDR < 0.05).

**Additional file 12: Table S16.** Primers and cDNA clones used for RNAi and in situ hybridization.

