## Supplementary material for "Lineage-specific, fast-evolving GATA-like gene regulates zygotic gene activation to promote endoderm specification and pattern formation in the Theridiidae spider": Addfile3 Figure S1-S10

### Additional file 3:
### Supplementary Figures
### (Figs S1 to S10)

**Lineage-specific, fast-evolving GATA-like gene regulates zygotic gene activation to promote endoderm specification and pattern formation in the Theridiidae spider**

Sawa Iwasaki-Yokozawa[1], Ryota Nanjo[1,2], Yasuko Akiyama-Oda[1,3,4], Hiroki Oda[1,2,*]
[1]Laboratory of Evolutionary Cell and Developmental Biology, JT Biohistory Research Hall, Takatsuki, Osaka 569-1125, Japan
[2]Department of Biological Sciences, Graduate School of Science, Osaka University
[3]PRESTO, Japan Science and Technology Agency, Kawaguchi, Saitama 332-0012, Japan
[4]Department of Microbiology and Infection Control, Faculty of Medicine, Osaka Medical and Pharmaceutical University

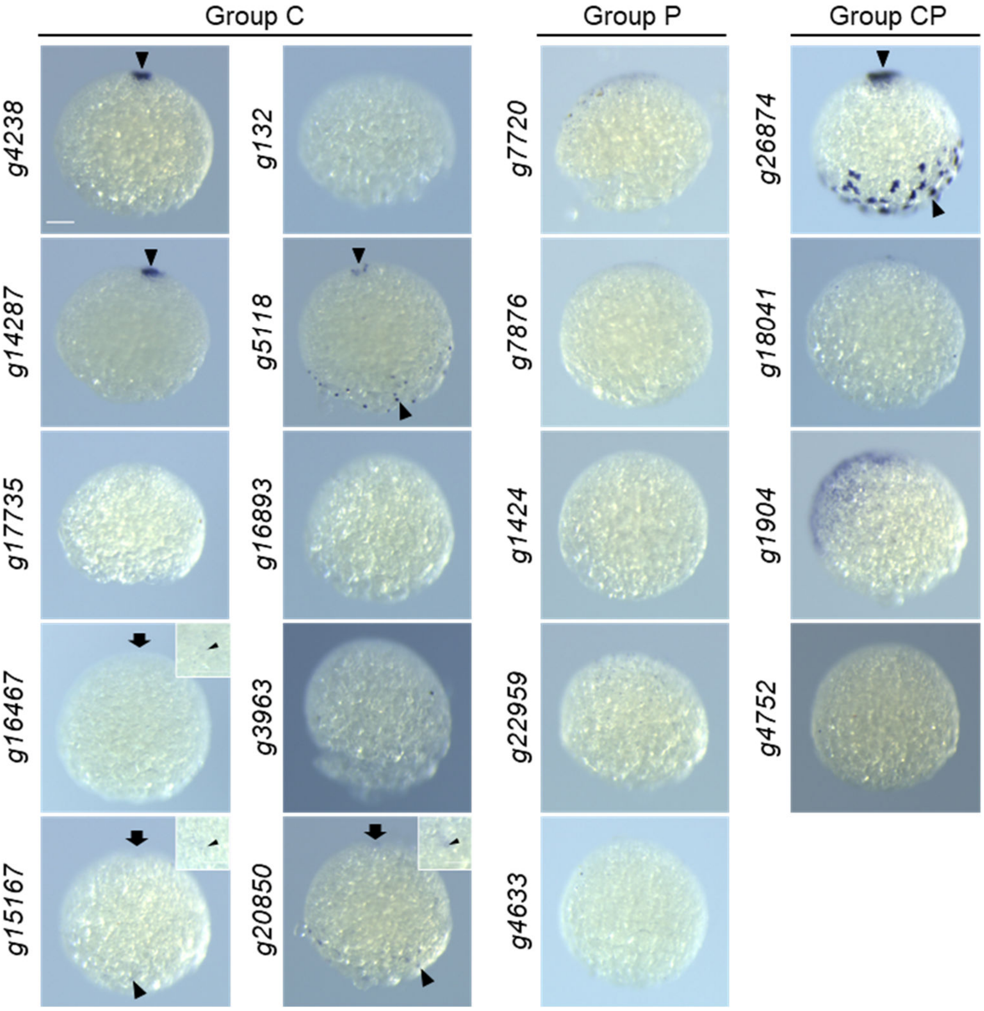


**Fig. S1. Chromogenic WISH of stage-3 embryos using probes for 19 selected DEG candidates.** For a pilot functional screen, 19 high-priority genes were selected from the DEGs identified by the comparative transcriptomes of cells isolated from 3 different regions of stage-3 embryo (see Fig. 2). The genes were grouped based on the DEG lists they come from (Groups C, P, and CP). All panels show the lateral view of stage-3 embryos stained by WISH using probes for the selected DEGs, as indicated. The forming germ disc is to the top. Transcripts for *g4238*, *g14287*, and *g16467* were detected at the embryonic pole, those for *g15267*, *g5118*, *g20850*, and *g26874* were both at the embryonic pole and in cells on the abembryonic side. Insets show high-magnification of a square region centered at the embryonic pole (fat arrows). Arrowheads denote reproducibly detected signals. Scale bar, 100 µm.

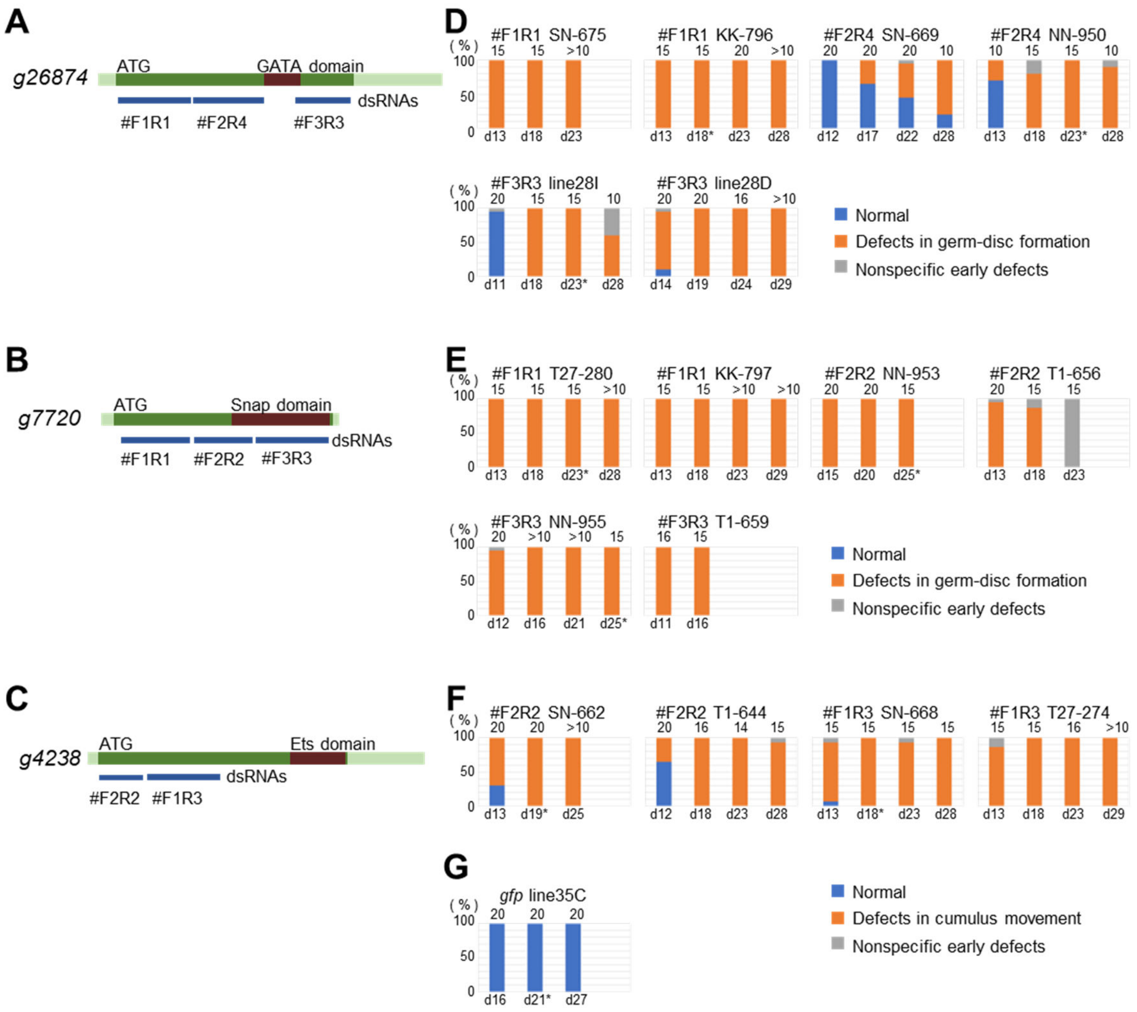


**Fig. S2. Validation of the specificity of the RNAi effects for *g26874*, *g7720*, and *g4238*. (A–C)** Schematic of the transcript for *g26874* (A), *g7720* (B), and *g4238* (C) and non-overlapping regions used for dsRNA synthesis (blue bars). **(D–G)** Phenotype expression in multiple egg sacs produced following injection of dsRNA for *g26874* (D), *g7720* (E), *g4238* (F), or *gfp* (G). Each graph shows data from one female injected with indicated dsRNA, and each vertical bar the percentages of embryos with indicated phenotypes in an egg sac, which was produced at indicated days after the first injection. The identification of individual females is also indicated (eg. SN-675). Number at the top of each bar is the total number of embryos examined. Development of 10 or more sibling embryos randomly selected from each egg sac was monitored using time-lapse microscopy, or with occasional visual inspection, under the stereomicroscope. The numbers of embryos that exhibited normal development (blue) and defects in the germ-disc formation or cumulus movement (orange) were counted. In certain egg sacs, some embryos that exhibited early defects prior to starting to form a germ disc, as denoted in gray. Because such defects sometimes occur regardless of RNAi, they were considered to be non-specific. Time-lapse movies of embryos from egg sacs indicated by asterisks have been deposited in a public repository [50].

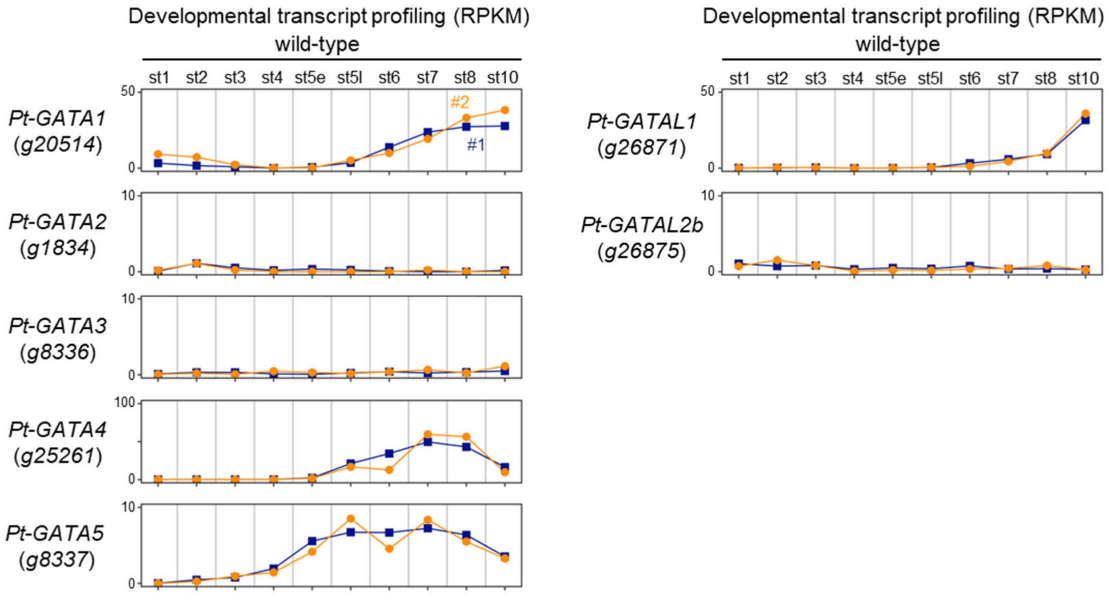


**Fig. S3.** Developmental transcript profiling of *P. tepidariorum* GATA family genes in wild-type embryos, based on public datasets (Iwasaki-Yokozawa et al., 2018). In addition to *fuchi* (g26874), there are seven annotated GATA family genes, *Pt-GATA1* (*g20514*), *Pt-GATA2* (*g1834*), *Pt-GATA3* (*g8336*), *Pt-GATA4* (*g25261*), *Pt-GATA5* (*g8337*), *Pt-GATAL1* (*g26871*), and *Pt-GATAL2b* (*g26875*). Two biological replicates are shown individually.

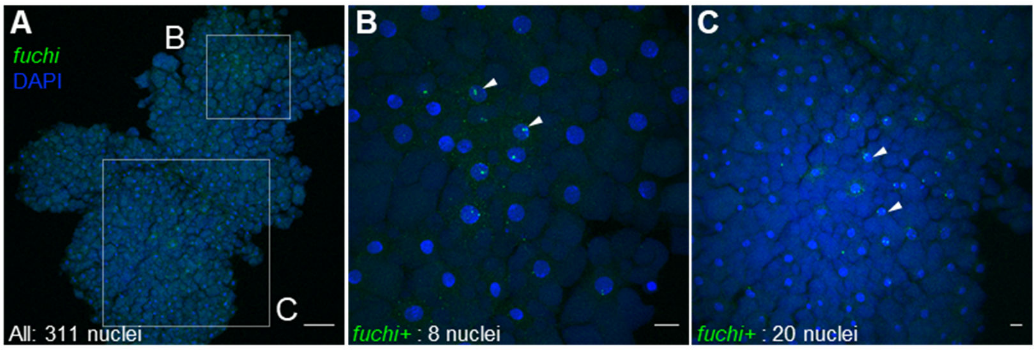


**Fig. S4. Cell counting in a stage-3 embryo stained for *fuchi*. (A–C)** A wild-type embryo at 15 h AEL was stained for *fuchi* transcript (green) and DNA (blue) using FISH, and the stained spherical embryo was flat-mounted with few parts lost. A total of 311 nuclei were observed (A). Around the embryonic pole (upper box in A), 8 nuclei were *fuchi* positive (magnified in B). On the ab-embryonic side (lower box in A), 20 nuclei were *fuchi* positive (magnified in C). Arrowheads point to examples of *fuchi*-positive nuclei. Scale bars, 100 µm in A; 20 µm in B, C.

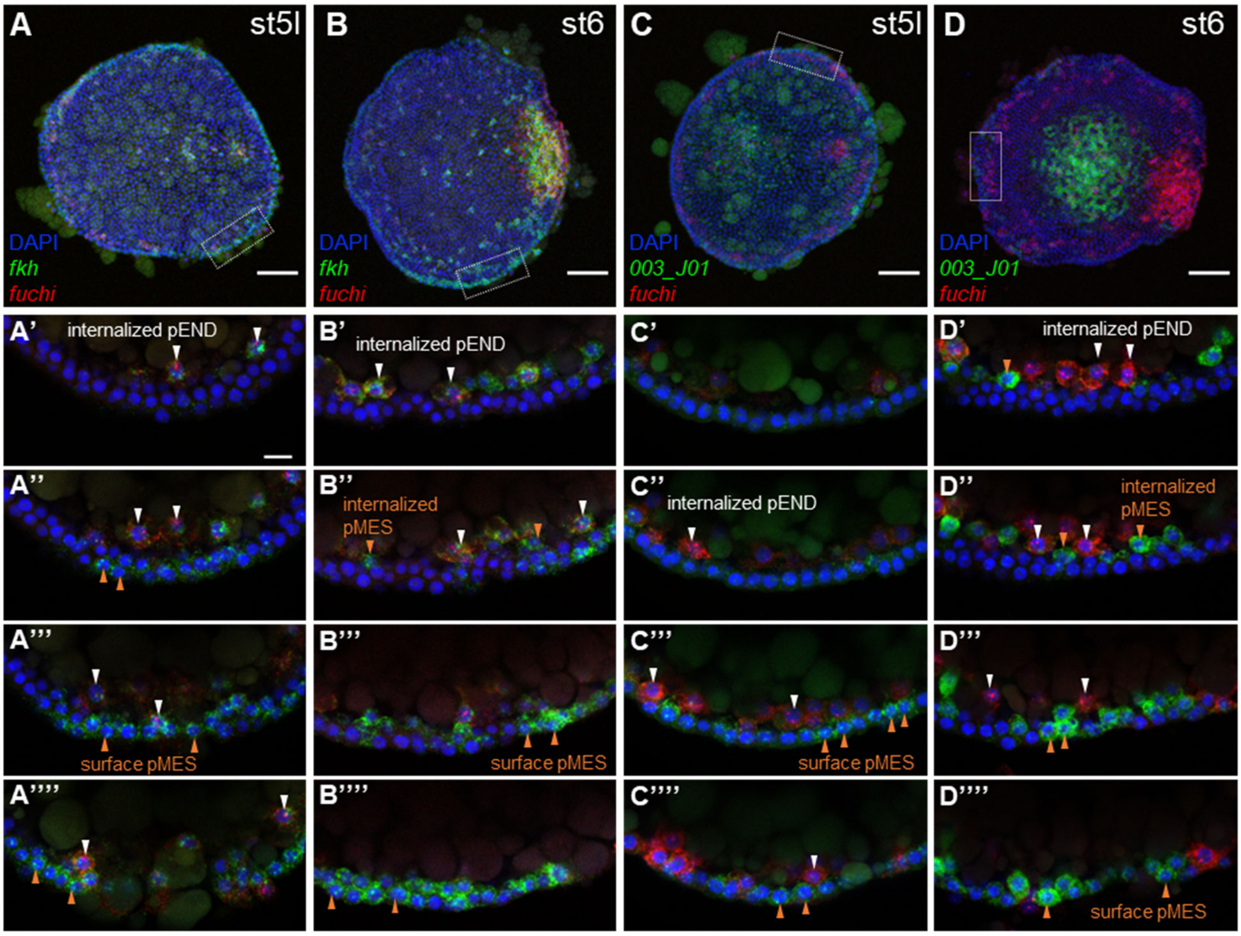


**Fig. S5. *fuchi* expression in pEND cells. (A–D)** Reconstruction of confocal stacks showing the top views of late stage-5 (A, C) and stage-6 (B, D) germ discs stained for *fuchi* (red), *Pt-fkh* (green in A and B; marker for mesoder plus endoderm) or *003_J01* (green in C and D; mesoderm marker) transcripts and DNA (blue). The areas boxed in A–D are magnified in the lower panels (A1-4, B1-4, C1-4, D1-4), where maximum-intensity projections of four successive sets of z-slices are shown; each projection image corresponds to 10-µm thickness. *fuchi* transcript was detected in *Pt-fkh*-positive cells that had been internalized earlier than *003_J01*-positive cells. These cells are pEND cells. Note that most, but not all, *003_J01*-positive cells on the surface and internalized are negative for *fuchi*, and that certain *Pt-fkh*-positive cells are negative for *fuchi*. Scale bars, 100 µm in A–D; 20 µm in A'.

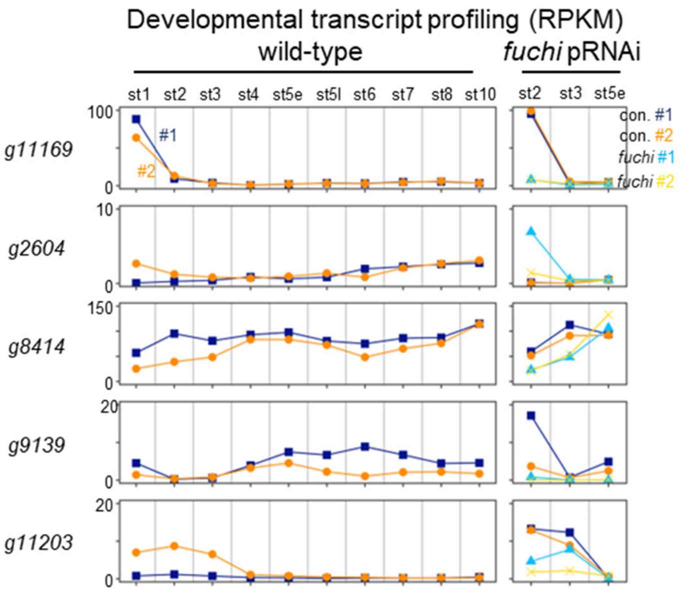


**Fig. S6. Characterization of five DEG candidates from the comparative transcriptome analysis of stage-2 *fuchi* pRNAi versus untreated embryos.** Graphs on the left side show developmental profiling of the transcript levels for the DEG candidates in wild-type embryos, based on public datasets [36], and those on the right side the effects of *fuchi* pRNAi on the transcript levels at the three stages examined in this study. In all graphs, two biological replicates of each sample type are shown individually.

93

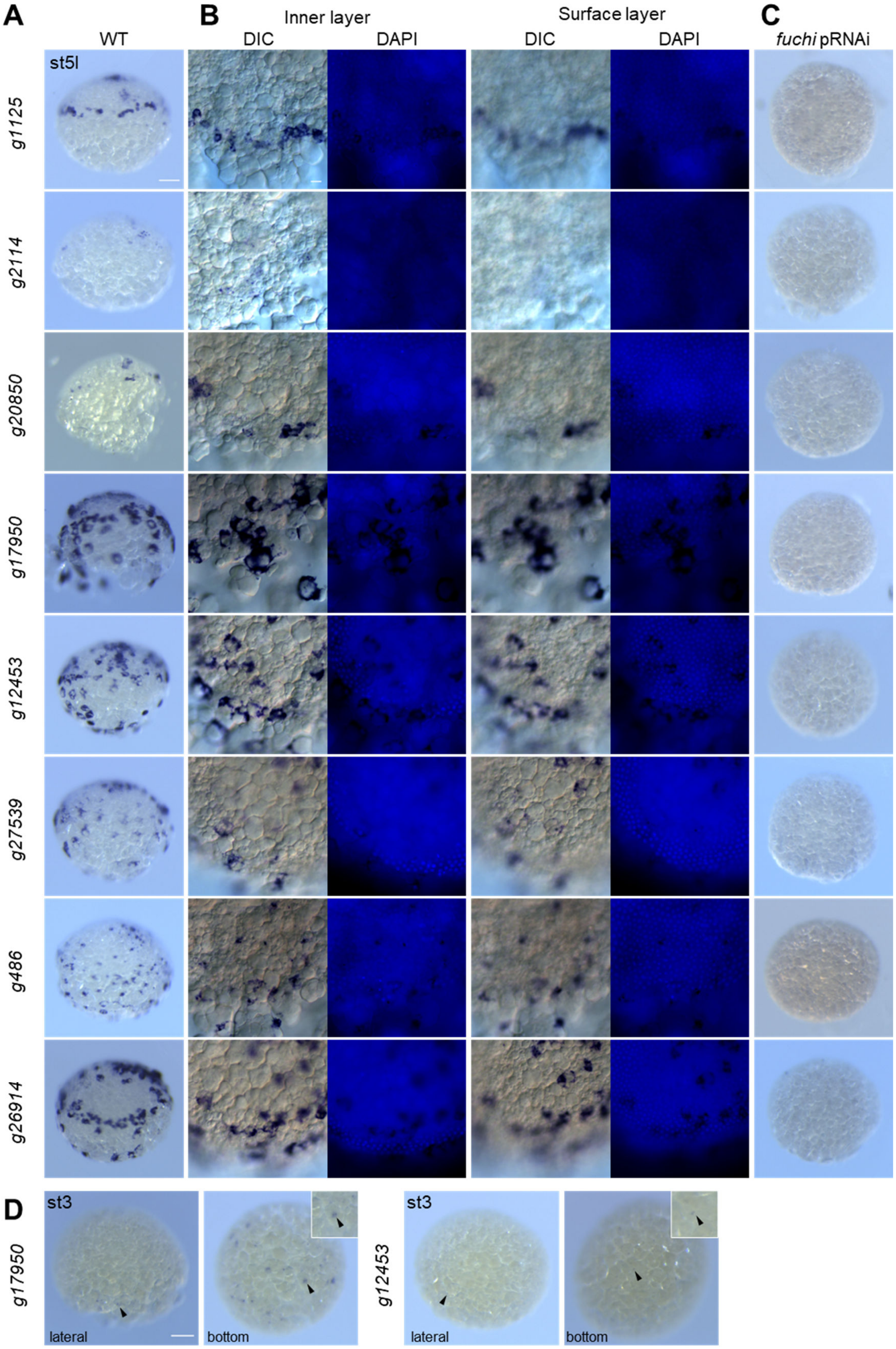
A
WT
B
Inner layer
DIC
DAPI
Surface layer
DIC
DAPI
C
fuchi pRNAi
st5l
g1125
g2114
g20850
g17950
g12453
g27539
g486
g26914
D
st3
g17950
lateral
bottom
st3
g12453
lateral
bottom

**Fig. S7. Characterization of the top-10 DEG candidates from the comparative transcriptome analysis of stage-3 *fuchi* pRNAi versus untreated embryos.** (A) Lateral view of stage-3 wild-type embryos stained by chromogenic WISH using probes for the DEG candidates indicated. The forming germ disc is to the top. No signals were obtained for two of the 10 DEG candidates (g*27086* and *g132*). (B) Close-up observation of germ-disc areas of the embryos shown in A, using differential interference contrast (DIC) optics. The focal plane for each field is set at two different depths; one plane is at an inner layer focusing on internalized pEND cells (left), while the other is at the surface layer (right). Counterstains with DAPI are also shown. (C) Lateral view of stage-3 *fuchi* pRNAi embryos stained in the same way as in A. (D) Lateral and bottom views of stage-3 wild-type embryos stained using probes for *g17950* and *g12453*. Arrowheads point to signals, and the boxed regions are magnified in insets. Stage-3 embryo stained for *g20850* has been shown in Fig. S1 (Additional file 3). Scale bars, 100 μm in A; 20 μm in B.

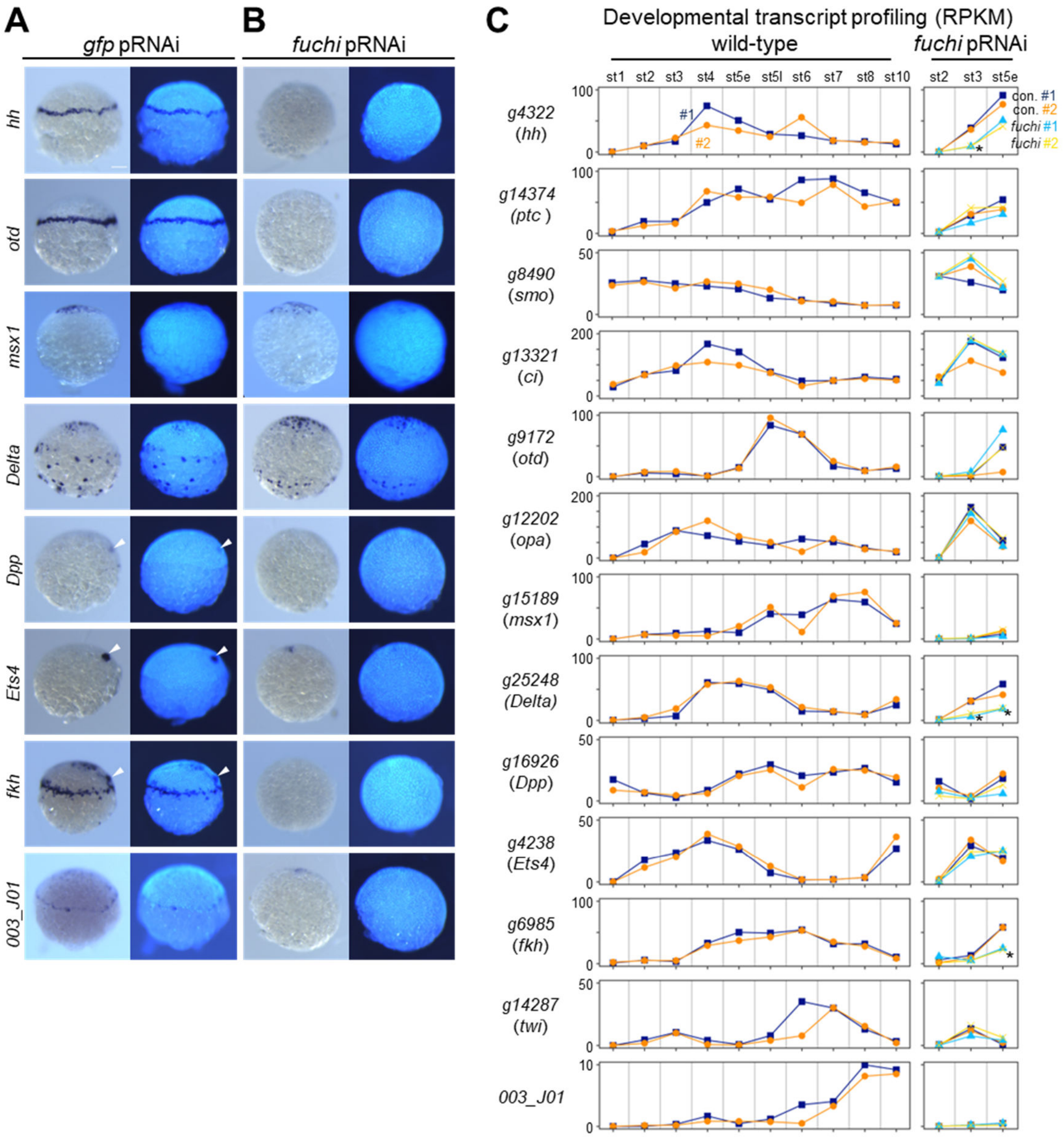


**Fig. S8. Effects of *fuchi* RNAi on expression of selected genes.** (A, B) Lateral view of late stage-5 *gfp* pRNAi (A) and *fuchi* pRNAi (B) embryos stained by chromogenic WISH using probes for the genes indicated. Counterstains with DAPI are also shown. The formed germ disc is to the top. White arrowheads indicate the migrating CM cells. Scale bar, 100 µm. (C) Graphs showing developmental profiling of the transcript levels (RPKM) for the indicated genes in wild-type embryos, based on public datasets [36], and the effects of *fuchi* pRNAi on the transcript levels at stage 2, stage 3, and early stage 5. Asterisks indicate that the corresponding comparative analysis identified the gene as a DEG (Additional file 9: Tables S10, S11). Two biological replicates are shown individually.

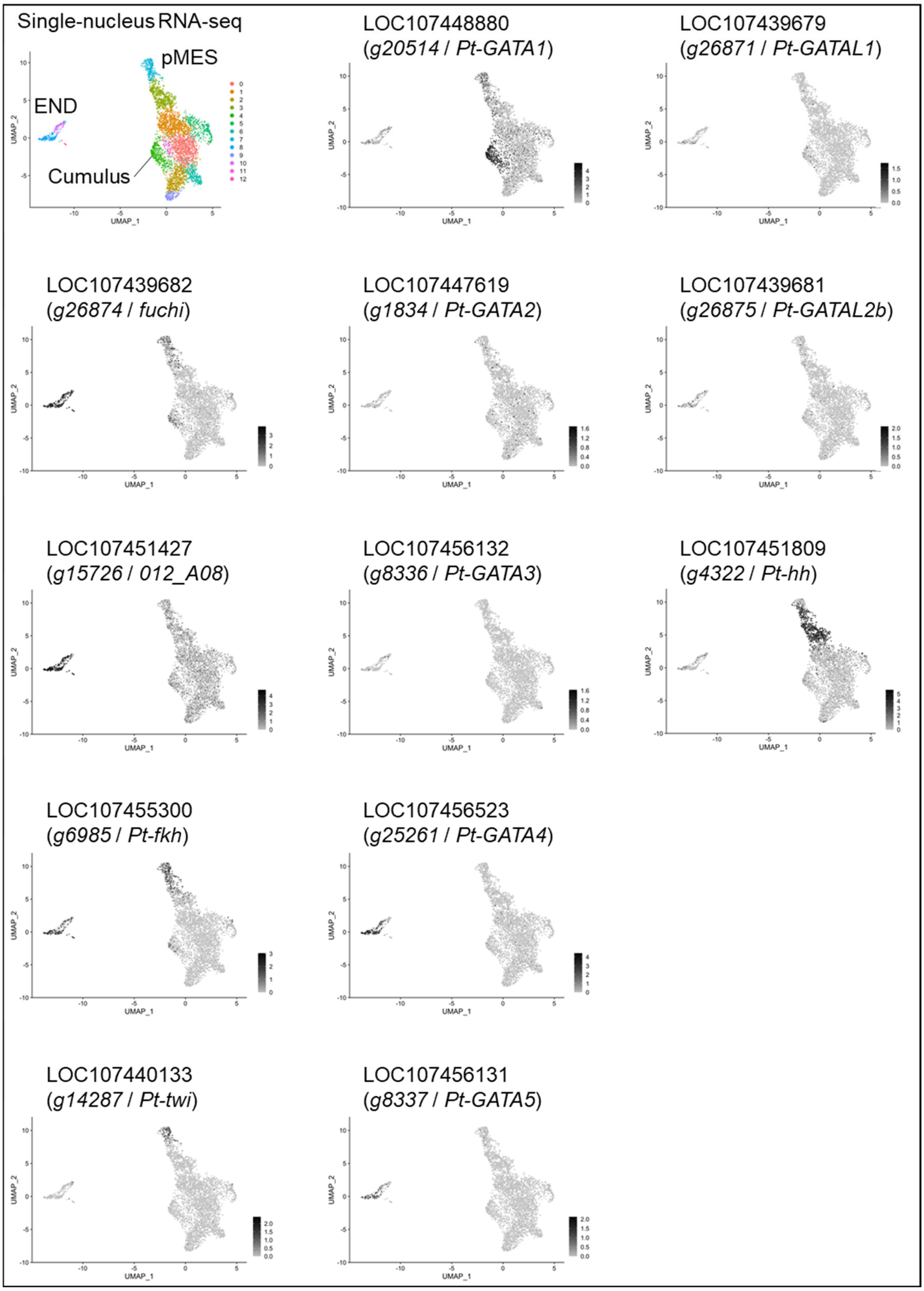


**Fig. S9. Single-nucleus transcriptome data showing endodermal and mesodermal cell populations in late stage-5 embryos.** Data shown are Uniform Manifold and Projection plots from clustering analysis of the single-nucleus transcriptome of *P.*

*tepidariorum* late stage 5 embryos described in a separate study [60,100]. Cell types are annotated in the top left plot, and expression of *fuchi*, known endodermal and mesodermal marker genes (*012_A08*, *Pt-fkh*, and *Pt-twi*), other GATA family genes (*Pt-GATA1* to *Pt-GATA5*, *Pt-GATAL1*, *Pt-GATAL2b*), and *Pt-hh* are represented in the other plots. The code prefixed with "LOC" indicates the gene identification used in NCBI.

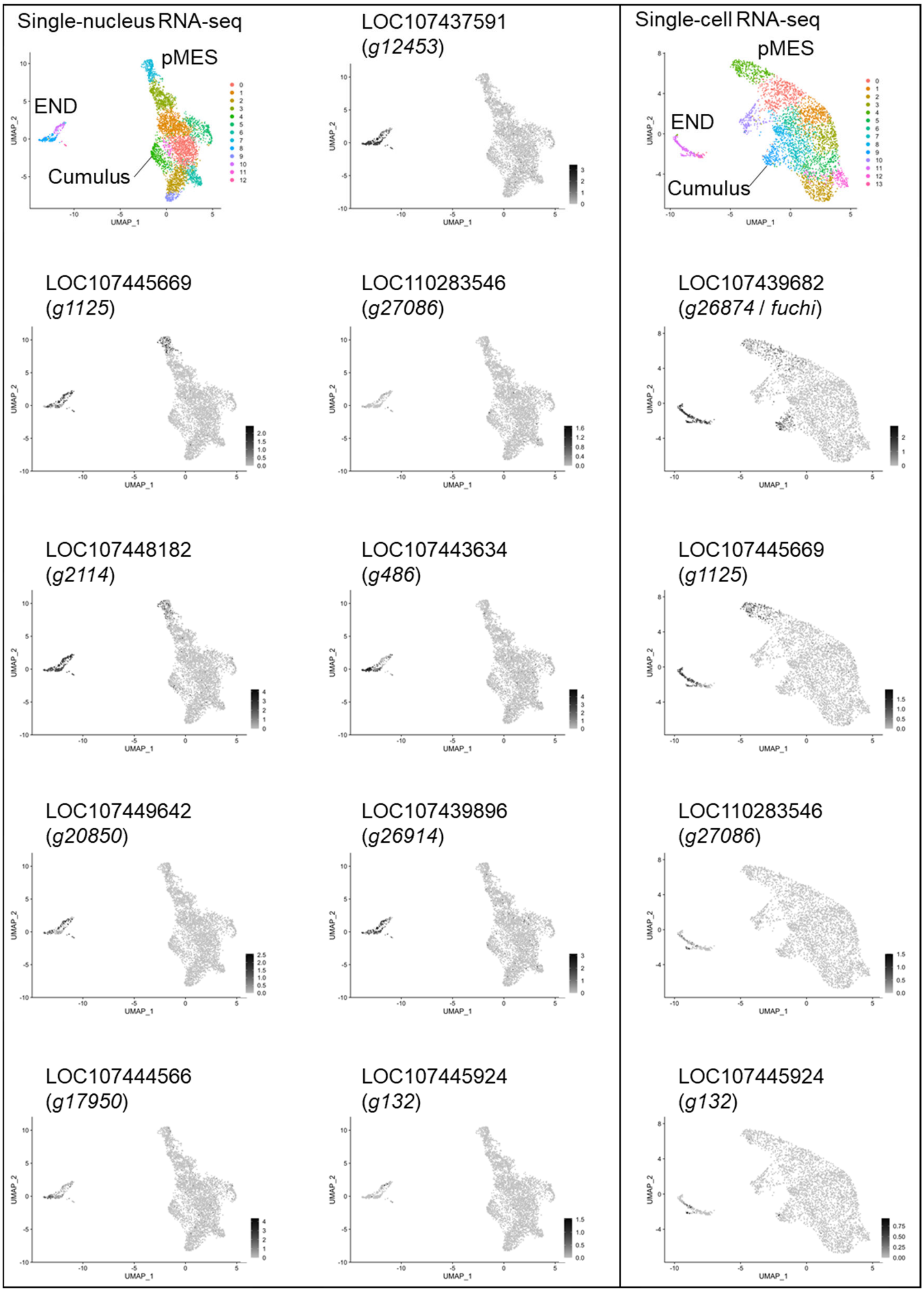


**Fig. S10. Single-nucleus and single-cell transcriptome data showing cell populations expressing the genes regulated by *fuchi*.** (A, B) Data shown are Uniform Manifold and Projection plots from clustering analysis of the single-nucleus (A) and single-cell (B) transcriptomes of *P. tepidariorum* late stage 5 embryos described in a separate study [60,100]. Cell types are annotated in the top left plot, and expression of the top-10 DEG candidates from the comparative transcriptome analysis of stage-3 fuchi pRNAi versus untreated embryos is represented in the other plots. The code prefixed with "LOC" indicates the gene identification used in NCBI.
